# Effects of habitat and fruit scent on the interactions between short-tailed fruit bats and *Piper* plants

**DOI:** 10.1101/2023.10.11.561911

**Authors:** Sneha Sil, Flo Visconti, Gloriana Chaverri, Sharlene E. Santana

**Author notes:** corresponding author: Sneha Sil **Email:**.

## Abstract

*Piper* is a mega-diverse genus of pioneer plants that contributes to the maintenance and regeneration of tropical forests. With deforestation and climate change threatening forest ecosystems, understanding the mutualism between *Piper* and its seed dispersers becomes especially important. In the Neotropics, *Carollia* bats use olfaction to forage for *Piper* fruit and are a main disperser of *Piper* seeds via consumption and subsequent defecation during flight. In return, *Piper* fruits provide essential nutrients for *Carollia* year-round. There is evidence that the types and diversity of *Piper* frugivores are influenced by the primary habitat of different *Piper* species (forest, gap), with forest *Piper* depending more on bats for seed dispersal; however, this pattern has not been tested broadly. We aimed to characterize and compare the interactions between *Carollia* and *Piper* across forested and gap habitats, and further investigate whether differences in fruit traits relevant to bat foraging (i.e., scent) could underlie differences in *Carollia-Piper* interactions. We collected nightly acoustic ultrasonic recordings and 24h camera trap data in La Selva, Costa Rica across 12 species of *Piper* (6 forest, 6 gap) and integrated this information with data on *Carollia* diet and *Piper* fruit scent. Merging biomonitoring modalities allowed us to characterize ecological interactions in a hierarchical manner: from general activity and presence of bats, to visitations and inspections of plants, to acquisition and consumption of fruits. We found significant differences in *Carollia-Piper* interactions between forested and gap habitats; however, the type of biomonitoring modality (camera trap, acoustics, diet) influenced our ability to detect these differences. Forest *Piper* were exclusively visited by bats, whereas gap *Piper* had a more diverse suite of frugivores; the annual diet of *Carollia*, however, is dominated by gap *Piper* since these plants produce fruit year-round. We found evidence that fruit scent composition significantly differs between forest and gap *Piper*, which highlights the possibility that bats could be using chemical cues to differentially forage for gap versus forest *Piper*. By integrating studies of *Piper* fruit scent, plant visitation patterns, and *Carollia* diet composition, we paint a clearer picture of the ecological interactions between *Piper* and *Carollia*, and plant-animal mutualisms more generally.

## Introduction

The interactions between plants and animals are crucial both for the ecology and evolution of species, and are responsible for maintaining and rebuilding healthy ecosystems (Whelan et al. 2008; Kunz et al. 2011). Bats, the only flying mammals, are particularly important in tropical and subtropical regions for the pollination and seed dispersal of hundreds of plant species, forming intricate networks mediated by morphological and behavioral co-adaptations (Mello et al. 2019). In the neotropics, the mutualism between two highly abundant and widespread taxa–short-tailed fruit bats (*Carollia* spp.; 9 species) and *Piper* plants (*Piper* spp.; ∼1200 neotropical species)–is an example of such a relationship. Via fruit consumption and subsequent defecation of seeds, *Carollia* disperse early, mid, and late succession *Piper* species, henceforth mitigating the changes to populations and community structure caused by deforestation and other forms of habitat alteration in tropical environments (Jones et al. 2009). In turn, *Piper* fruits make up to 35% of *Carollia*’s annual diet (Santana et al. 2021; Lopez and Vaughan, 2007; Maynard et al. 2019) and provide a consistent source of nutrients for these bats (Fleming 1991; Gelambi and Whitehead, 2023). As a result, *Piper* has likely evolved fruit structures, shapes, textures, and scents ideal for bats to locate and feed on them (Hodgkison et al. 2013; von Helversen and von Helversen 1999; von Helversen, Holderied, and von Helversen 2003; Simon et al. 2011), and in turn *Carollia* has evolved a specialized sense of olfaction, chemical preferences, and foraging behaviors that presumably aid in locating ripe *Piper* fruits (Thies and Kalko 2004; Santana et al. 2021; Yohe et al. 2021). Furthermore, behavioral experiments have shown *Carollia* primarily utilizes olfaction to locate fruiting patches and then echolocation when closer to their target before snagging fruit, and these bats only seem to attempt consumption of *Piper* fruits when appropriate scent cues are present (Thies et al. 1998; Leiser-Miller et al. 2020).

Most Neotropical *Piper* plants produce green infructescences with small seeds and a distinctive bouquet of volatile organic compounds (VOCs) when ripe (Thies and Kalko, 2004; Santana et al. 2021). These VOCs can be secondary metabolites produced during ripening that can act as signals adapted to target mutualistic frugivores, and include various compounds such as terpenes, alcohols, and carbonyl compounds (Santana et al. 2021). Santana et al. found that bats prefer samples enriched with the *Piper* VOCs 2-heptanol and alpha-caryophyllene, indicating that these compounds could have a role in attracting bats to ripe *Piper* fruits (Santana et al. 2021). An aspect that remains unknown, however, is the extent to which fruit ripeness and the strength of the chemical signal generated by its scent may influence bat foraging behavior, including how frequently bats visit different *Piper* species. For example, *Piper* species with strong scent signals or VOCs preferred by bats might be more readily visited and consumed by bats than those that do not, in which case bats may spend more time inspecting fruits and flying around the plant in general.

While the *Carollia-Piper* mutualism has been characterized on many fronts, the patterns of interactions between these bats and plants across habitats have received less attention. This information is critical for understanding how dynamic these interactions are across space, the role of these species in local ecological communities, and their importance in ecosystem resilience. At one Panamanian site, Thies and Kalko found that *Piper* species differed in their time of ripening and seed disperser spectrum, and thereby provided the broad characterization of two major *Piper* ecotypes: “forest” *Piper*, which exhibit short and staggered fruiting peaks, fruits that ripen in the evening, and a narrow spectrum of frugivores (bats; *C. castanea, C. perspicillata*), and “gap” *Piper* with extended fruiting seasons, fruits that ripen early in the morning, and a larger range of seed dispersers (bats, birds, insects) (Thies and Kalko 2004).

While they posited that the differences in flowering phenology between forest and gap *Piper* were primarily caused by abiotic factors, they suggested differences in fruiting patterns (staggered vs. continuous; morning vs. evening) were influenced by the respective spectrums of seed dispersers across each habitat. That is, the long and overlapping fruiting periods of gap *Piper* species would be associated with a larger spectrum of dispersers that would possibly mitigate the challenges of seed dispersal into spatially unpredictable germination sites (Thies and Kalko 2004). Despite this landmark study, however, it is not known whether the forest and gap *Piper* ecotypes are generalizable to other *Piper* species and sites in the Neotropics. Here, we aim to help fill this knowledge gap by contrasting frugivore visitation patterns to different *Piper* species in Costa Rica; we relate our results to the *Piper* habitat-based ecotypes defined by Thies and Kalko and to fruit traits (i.e. fruit scent) known to play an important role in this mutualism. We hypothesize that habitat plays a role in defining the community of frugivores that feed from *Piper* plants, and predict that there will be a greater diversity of frugivores visiting gap *Piper* species compared to forest *Piper*, with the latter being consumed exclusively by bats (consistent with the Thies and Kalko 2004 study). We also hypothesize that differences in fruit scent VOCs between forest and gap *Piper* may contribute to differences in how attractive they are to bats, and hence influence bat visitation and consumption patterns across habitats.

Researchers traditionally characterize plant-bat interactions by acoustic monitoring and mist-netting of live animals in the field for diet studies (Fraser et al. 2020). Here, we applied an integrative approach that provides a more nuanced “plant perspective” to documenting these interactions. We used three biomonitoring methods –camera trap videos, ultrasonic acoustic recordings, and dietary analyses– for a detailed comparison of the *Carollia-Piper* mutualism across forest and gap habitats, and combined our results with information about *Piper* fruit chemical signals relevant to bat consumption patterns. Working in a Costa Rican site, we evaluated the visitation frequency of bats and other frugivores to *Piper* plants in forest and gap habitats via nightly ultrasonic acoustic recordings and 24h camera traps and complemented these data with our published data on *Piper* consumption by *Carollia* and *Piper* fruit VOCs, all collected at the same site. We find that this approach allows us to characterize ecological interactions in a hierarchical manner: from general activity and presence of bats, to visitations and inspections of plants, to acquisition and consumption of fruits. Our study thereby provides a novel interpretation of the *Carollia*-*Piper* mutualism and provides insight into how best to document the interactions within this important ecological system.

## Methodology

### Study site

This study was conducted at the Organization for Tropical Studies’ La Selva Biological Reserve, Costa Rica (herein La Selva). The reserve comprises 1,600 ha of protected area spanning primary premontane and tropical wet forest, secondary forest, and abandoned agricultural land. *Piper* is highly diverse at La Selva, with over 50 recognized species (OTS 2023), which can be roughly classified into the “gap” (early-succession) or “forest” (mid-to late-succession) categories of Thies and Kalko (2004) (see Supplemental Information Table S1; Grieg 1993). Three *Carollia* species (Chiroptera: Phyllostomidae) occur at La Selva (*C. castanea*, 11 g; *C. sowelli*, 18 g; and *C. perspicillata*, 21 g; Santana et al. 2021; Fig. 1c); these are some of the most abundant bats at the site year-round and coexist with about 62 other bat species (OTS 2023). This research was conducted under Costa Rican permit SINAC-ACC-PI-R-107-2019. All procedures were approved by the Institutional Animal Care and Use Committee of the University of Washington, Seattle, USA (protocol #4307-02).

**Figure 1:**
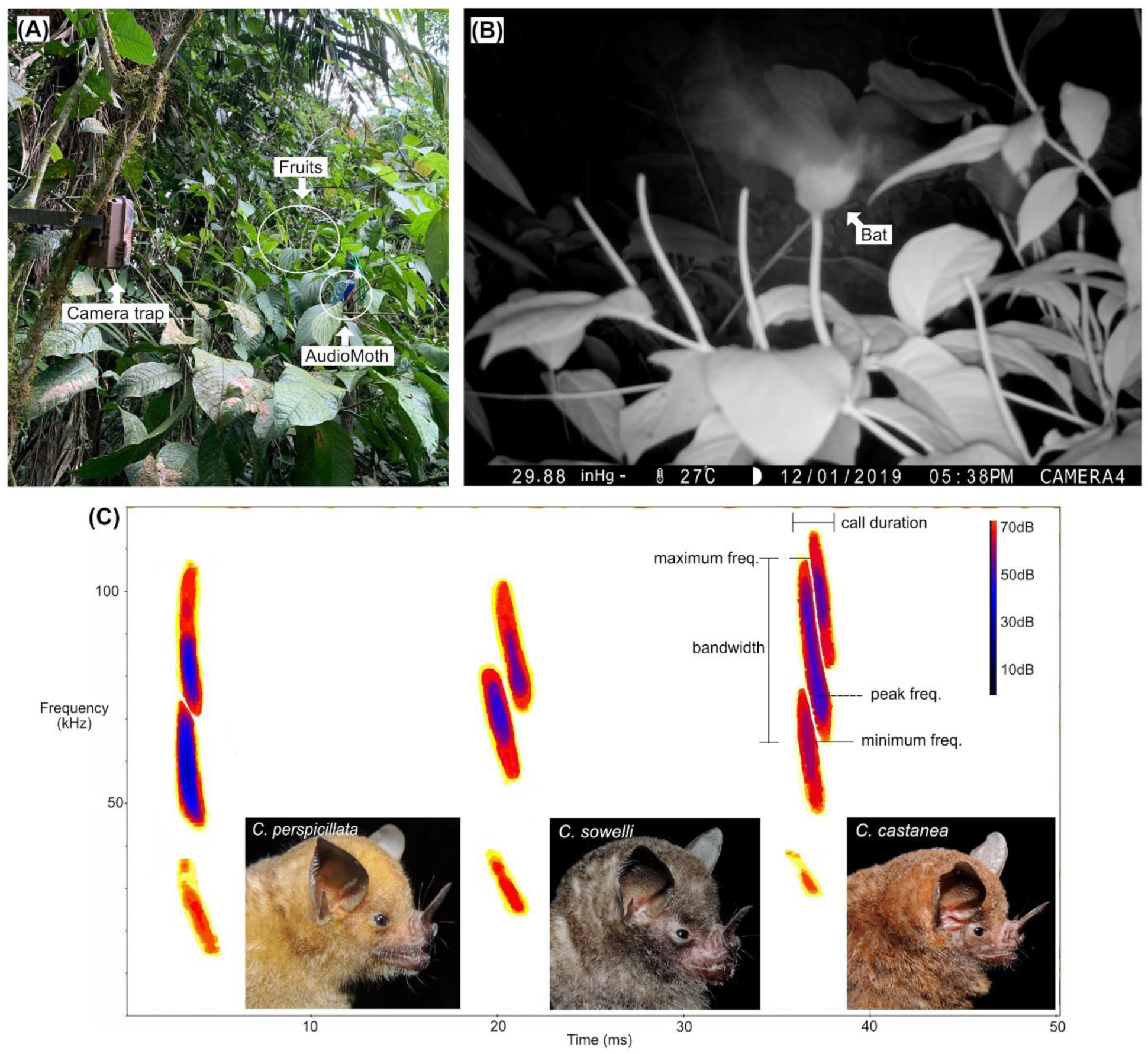
Experimental setup including camera trap and ultrasonic acoustic recorder (AudioMoth) deployed at a *Piper sancti-felicis* plant in the field (A), a video frame showing a bat collecting a fruit at the same plant (B; video available as a supplemental file [Supplemental Video 1]), and reference echolocation calls for *Carollia perspicillata, C. sowelli,* and *C. castanea* (C; spectrograms generated in BatSound v4.4). Analysis of acoustic data was performed using the parameters marked in the spectrogram (call duration, peak frequency, minimum frequency, maximum frequency, and bandwidth of the main harmonic; see Table S2). *Carollia* photos credit: David Villalobos Chaves.

### Recording setup

We documented bat activity and behavior at 45 plants across 12 species of *Piper* (6 forest, 6 gap; Table 1) for 1-211 days per plant between 2019 and 2021 (Table S1). We selected *Piper* plants on the basis of three criteria: (1) plants had at least one fully formed (presumed ripe or close-to-ripe) fruit; the fruits of most neotropical *Piper* species remain a shade of green when ripe but become noticeably plump and softer when they approach ripeness; (2) fruits were accessible to place acoustic recorders and cameras no more than 50 cm (acoustics) or 5 m (cameras) away from fruits (Fig. 1a); (3) plant location maximized spatial distance among plants of the same species (at least 3 m, but typically tens to hundreds of meters apart; Fig. 2, Table S1). For video documentation of frugivores at *Piper* plants, we used motion-activated BrowningAdvantage Spec Ops Full HD Video Trail Cameras (Browning Trail cameras, USA; Model BTC-8A), which were strapped to trees, lianas, poles, rails, or other available structures and positioned to ensure the fruits were centered within the field of view (Fig. 1a). Cameras were set to capture HD videos at a 1920 x 1080, 60 fps resolution, with motion detection at a minimum of 60 ft. and a trigger speed of 0.4 sec. The cameras recorded for 24h each day, using an infrared function during the night, and set to record for 20 seconds as soon as movement was detected. Sequential 20 second videos were stored when movement was detected for longer periods of time. For acoustic documentation of bats during the night, we used AudioMoths (Open Acoustic Devices), which are full-spectrum acoustic loggers based on the Gecko processor range from Silicon Labs. We placed these close to fruits (≤50 cm), encased in the AudioMoth IPX7 Waterproof Case. We set AudioMoths to record starting at dusk and to span the known high activity period of *Carollia* (5-8 PM local time), using a sample rate of 256 kHz, medium gain, for 10 second intervals every 20 seconds. We chose these settings to increase our chances of detecting *Carollia’s* relatively “quiet” echolocation calls, and produce a manageable amount of data, respectively. We monitored plants every 2-3 days and stopped video and audio recordings as soon as the focal fruit(s) had been removed from the plant, and no plants were recorded more than once. A few plants, however, were video recorded for a much longer time because we had to leave cameras deployed and unattended during lockdowns and travel restrictions associated with the COVID-19 pandemic.

**Figure 2.**
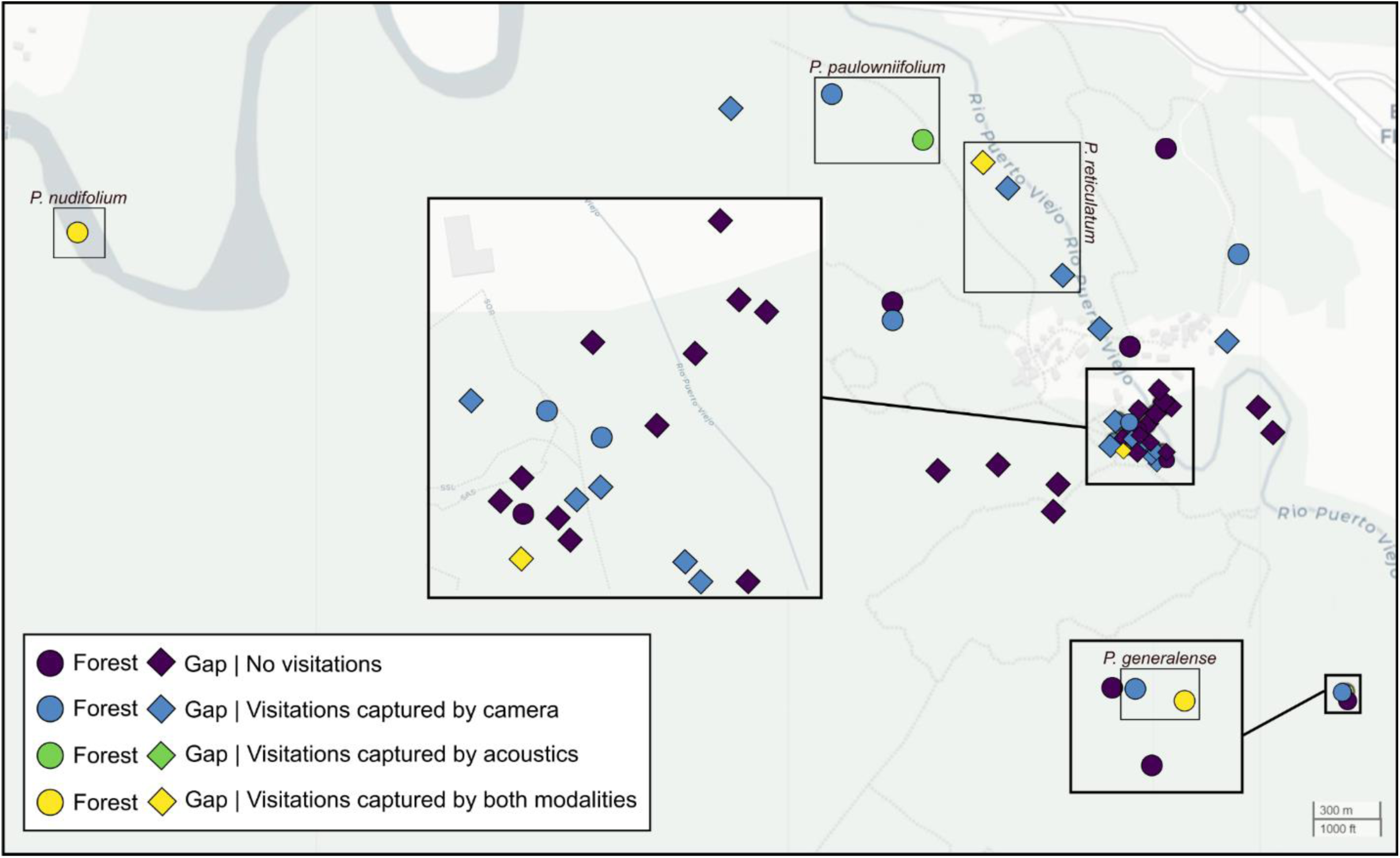
Map of the study area at La Selva Biological Reserve, Costa Rica, showing the locations of all *Piper* plants, within forest and gap habitats, where camera traps and acoustic recorders were deployed. Each plant is color-coded based on whether *Carollia* visitations occurred and how these visitations were documented: by camera traps, acoustic recorders, or both. Groups of *Piper* species showing activity by both camera traps and acoustic recordings are labeled as well.

**Table 1.**
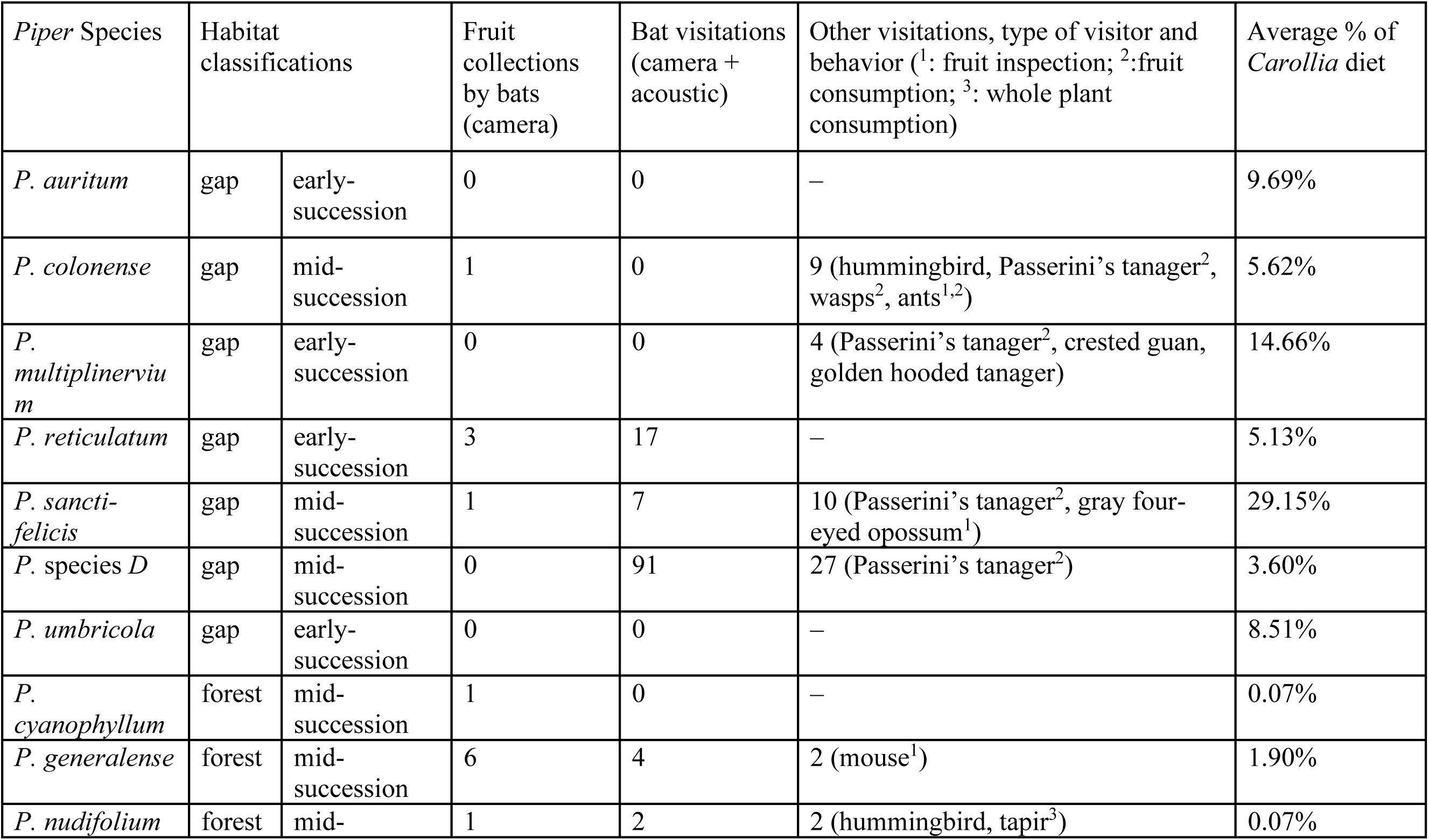
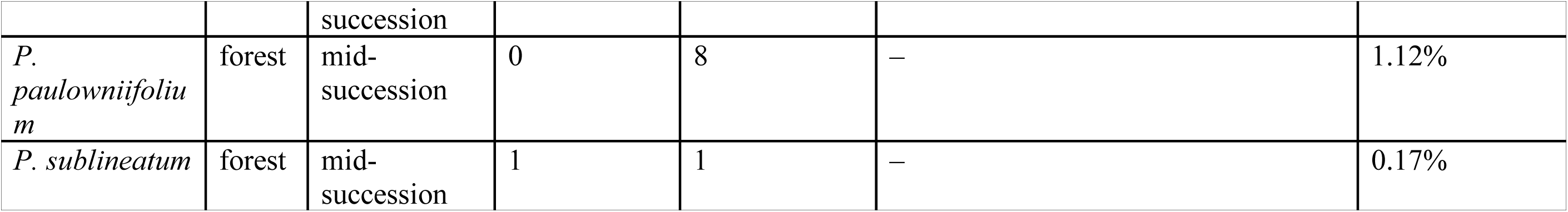
The 12 *Piper* species at La Selva, Costa Rica, focal to this study, their habitat classification, number of fruit collections and visitation events by bats and other frugivores recorded by camera traps, and the average percent of each species in the annual diet of *Carollia sowelli, C. perspicillata,* and *C. castanea* (from the literature, see text for sources).

### Camera trap video analysis

One of us (F.V.) performed video analysis to avoid bias in the results. We analyzed videos collected from camera traps using QuickTime Player 8 on a macOS operating system, and took note of: the organism(s) observed in the recording to the lowest possible taxonomic level (e.g., bat, tanager, tapir, spider, rodent, and so on), the action performed by the organism, and the time and date at which this behavior took place. We first observed each 20 second video at normal speed playback to help identify the source of movement, since the camera trap sensor was sometimes triggered by leaves or branches being blown by wind. When an animal was encountered in the videos, we would play the video again at half speed at least once or twice to determine what behavior was being performed. Bats circling plants move at a fast speed; therefore, many videos had to be analyzed two or three additional times at half speed to properly identify behavior. Additionally, we analyzed videos about 4-5 times at half speed and original speed if they contained activity from more than one animal, such as multiple tanagers, so we could accurately take notes on each individual’s behavior. We performed classification of animals that were not bats with the aid of field guides containing physical descriptions and images of the different animal species found across Costa Rica (Garrigues and Dean 2007).

### Acoustic analysis

To create quantitative and qualitative references for manual *Carollia* echolocation call identification in our field data, we compiled a call library of search-phase echolocation call recordings of *C. perspicillata, C. sowelli,* and *C. castanea*. These calls were recorded with a condenser microphone (microphone capsule CM16, CMPA preamplifier unit, Avisoft Bioacoustics, Berlin, Germany). We generated spectrograms (e.g. Fig. 1c) using RavenPro v. 1.6.2 (512 FFT Hanning window; 95% overlap; K. Lisa Yang Center for Conservation Bioacoustics at the Cornell Lab of Ornithology 2022), and collected the following parameters to act as a quantitative reference: call duration (ms), 90% call duration (ms), minimum frequency (kHz), maximum frequency (kHz), peak frequency (kHz), 95% frequency (kHz), delta frequency (kHz), and 90% bandwidth (kHz) (Table S2). Call duration, peak frequency, and delta frequency are widely used to characterize echolocation vocalizations (Luo et al. 2019), and the general frequency ranges and shape of echolocation calls are indeed relevant for qualitative manual ID. However, considering the lack of published *Carollia* spp. call library data, we collected extra parameters to increase the reliability of our manual ID method and to serve for future reference (Table S2). This preliminary step of analyzing focal call data and creating call guides is essential for proper acoustic identification, as bat calls may be only accurately identified by known qualitative and/or quantitative measures (Fraser et al. 2020). However, classification of *Carollia* calls down to the lowest taxonomic level is unreliable due to both the intraspecific and interspecific variability in type of call (social, search-phase), diagnostic call traits, and similar call structure to other Phyllostomidae species (Obrist 1995, Barclay 1999, Russo et al. 2018, Fraser et al. 2020, Fig. 4.5 in Collen 2012). Identification to the species level is especially difficult for low-duty cycle call species such as *Carollia* spp., because their calls exhibit the most intraspecific and intraindividual flexibility associated with different tasks and habitat effects (Russo et al. 2018). Our focal call parameters showed significant overlap between *C. castanea, C. perspicillata,* and *C. sowelli* echolocation calls (Fig. 1c); therefore, we aimed to mainly identify calls to the *Carollia* genus when possible.

The main challenge in analyzing passive acoustic recordings from a tropical forest site is environmental clutter: humidity, vegetation and foliage, and other animal sounds can cause echoes and additional noise into the path of the incoming sound (Fraser et al. 2020). These factors are unavoidable; as a result, our field data contained significant background noise. Additionally, we accrued a massive dataset which was impractical for one researcher to go through manually (characteristic of most experiments utilizing passive acoustic monitoring [Fraser et al. 2020]); therefore, we used a semi-automated method to sort through our large, noisy acoustic dataset. We developed a parallel-processing filtering program in MATLAB v. 9.12.0 (The MathWorks Inc. 2022; Fig. 3), which sorted through the dataset using a bandpass filter (butterworth) to filter out noise below the minimum frequency threshold of *Carollia* calls (approximately 45kHz, according to our focal parameters; Table S2). Then, the program generated a power spectrum (pwelch) used to filter the acoustic files into two categories: containing bat calls (above a threshold frequency) or mainly consisting of noise (below the threshold frequency). The threshold frequency, like the bandpass filter, was chosen based on the focal call data parameters (in this case, the peak frequencies of three *Carollia* species). If the peak frequency of the filtered signal was in the range of *Carollia* search-phase echolocation call peak frequency (anywhere from 60-80kHz depending on the species [Table S2]), this indicated high activity within that frequency and the likely presence of bats in the habitat where the calls were recorded. The algorithm ran on the dataset twice with two different bandpass and peak frequency threshold parameters; once with more sensitive parameters (type II error) and once with more specific parameters (type I error). The overall goal of this program was to sort through the large dataset and set aside a reasonable number of files (combined from the results of both algorithm runs [Table S3]) for a researcher trained on spectrogram analysis of *Carollia* focal search-phase echolocation calls (S.Sil) to analyze manually. We deemed this hybrid approach the best way to deal with the large dataset and noise present in the data considering that completely automated identification can generate significant error rates which could influence our characterization of *Carollia-Piper* interactions across habitats (Russo and Voigt, 2016; Rydell, Nyman, Eklof, Jones and Russo 2017; Barre et al. 2019).

**Figure 3.**
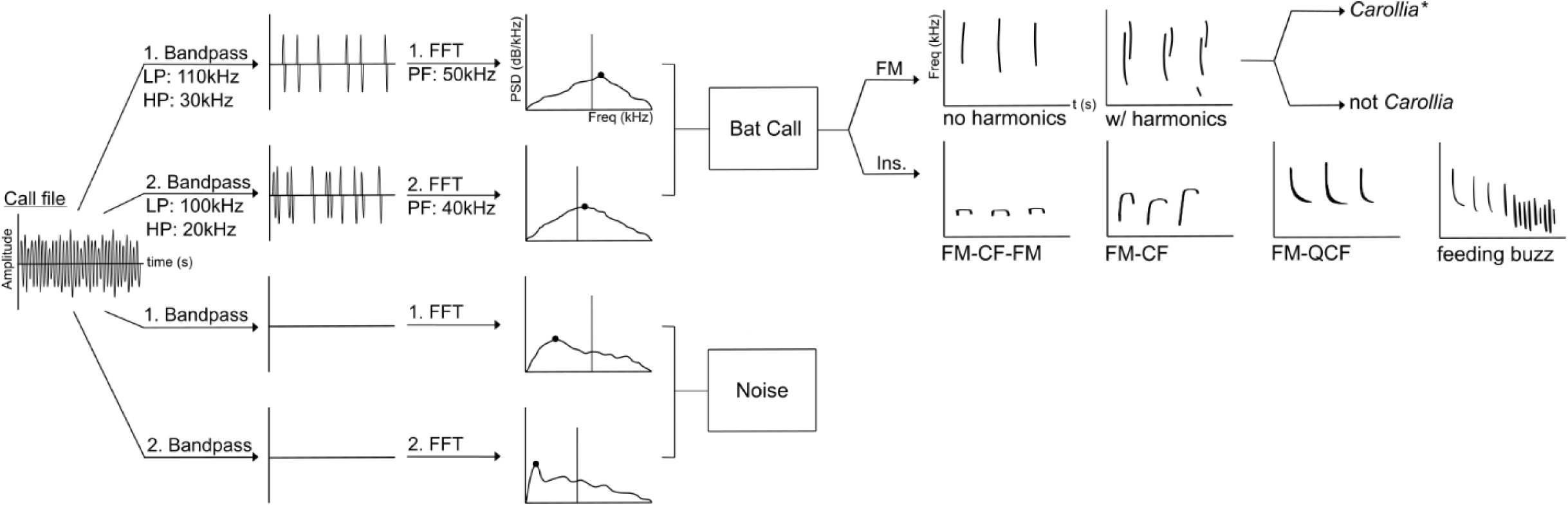
Visual representation of our algorithm developed to first filter (LP: low pass, HP: high pass) through the large acoustic dataset and identify files with bat calls present (PF: peak frequency), and the subsequent criteria used to manually categorize these files into various call types (FM: frequency modulated, Ins: insectivorous). The first run (LP: 110kHz, HP: 30kHz) settings were more specific, and the second run (LP: 100kHz, HP: 20kHz) settings were more sensitive. FM calls were split into calls with no harmonics and calls displaying harmonics (which were then qualitatively determined to be *Carollia* calls or not). Insectivorous calls were split into FM-CF-FM calls (frequency-modulated, constant-frequency, frequency-modulated), FM-CF calls (frequency-modulated, constant frequency), and FM-QCF (frequency-modulated, quasi-constant frequency) calls. Feeding buzzes were also noted. Results with the total numbers of each call type identified at each *Piper* plant analyzed after filtering can be found in Table S3. *See Figure 1c for criteria on qualitatively identifying *Carollia* bat calls.

Subsequently, one of us (S.Sil) carried out manual identification of bat calls across individual *Piper* plants and species to avoid bias in the results. We generated spectrograms to view calls using RavenPro v. 1.6.2 and BatSound v. 4.4 (512 FFT Hanning window, 95% overlap; K. Lisa Yang Center for Conservation Bioacoustics at the Cornell Lab of Ornithology, 2022; Pettersson Elektronik AB, 2016). To supplement the *Carollia* focal data we collected, we also used published Phyllostomidae search-phase echolocation calls as a guide (Fig. 4.5 from Collen, 2012). To avoid confusion amongst the cluttered environment and presence of other bat species, we only noted calls above 40kHz (Table S4; based on the typical minimum frequency of *Carollia* calls being approximately 45kHz [Table S2]). Additionally, we only identified *Carollia* calls as such if they matched our focal data, consisted of at least two harmonics, and had a high signal to noise ratio on the main harmonic.

**Figure 4.**
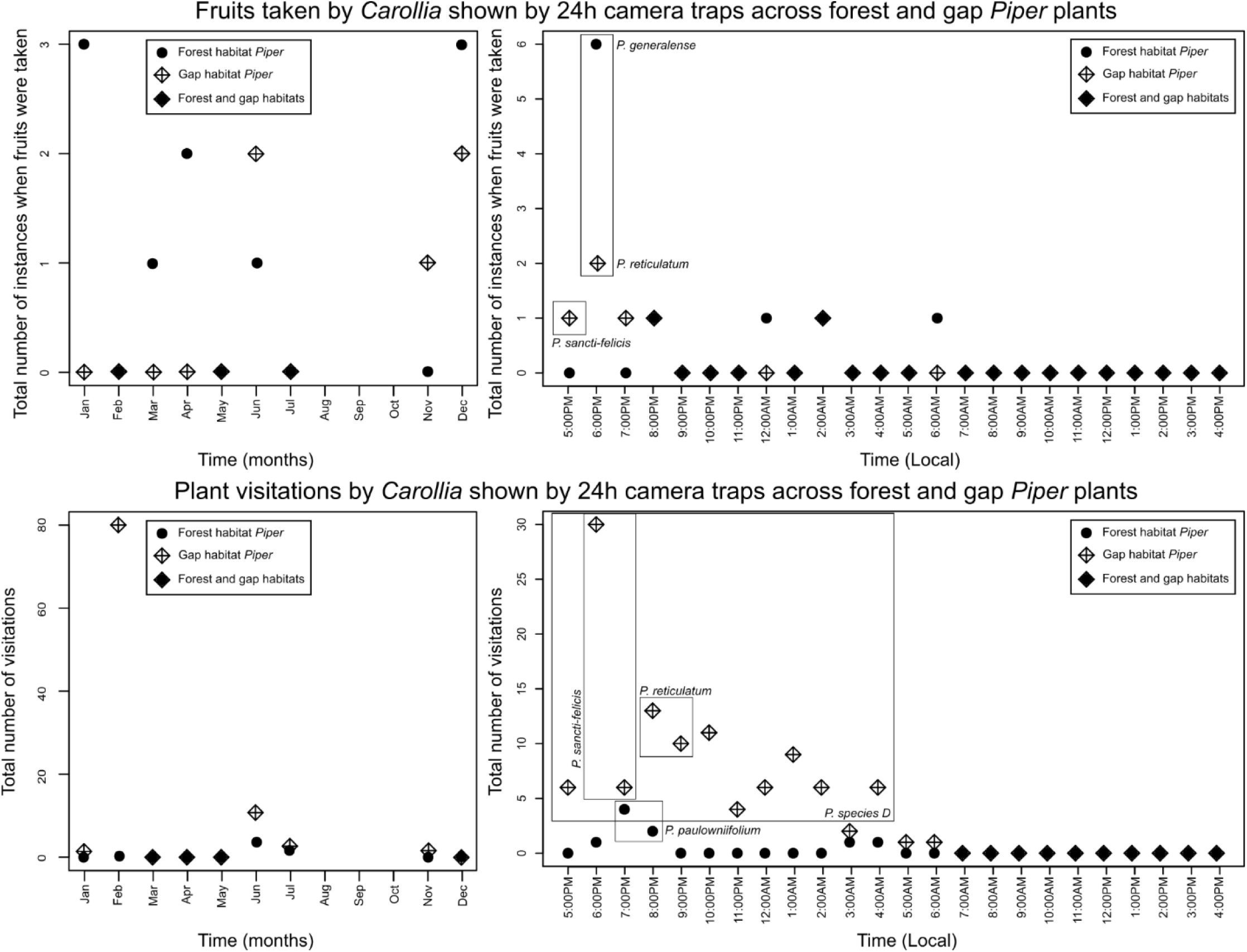
Temporal patterns of bat activity across forest and gap *Piper* plants. Top: Instances of bats taking fruits as shown by 24h camera traps across forest and gap *Piper* over the course of the year and throughout the day. Data are shown starting from sunset (5:00 PM local time), when bat foraging begins. Bottom: *Carollia* visitations to plants as shown by 24h camera traps across the same temporal scales. High activity peaks are noted on the plots with the *Piper* species at which they occurred. Data are the visitations and instances of fruit acquisitions added across all plants of a species for a given month/hour throughout the length of the study.

### Diet and fruit scent data

As a third proxy of *Carollia-Piper* interactions, we compiled the percentage of *Piper* species (33 documented to date; 24 forest, 9 gap habitat) found in the respective diets of *C. castanea, C. sowelli,* and *C. perspicillata* at La Selva. These data were based on fecal samples collected from hundreds of free-ranging bats at La Selva and published by one of us (Santana et al. 2021, which incorporated data from Lopez and Vaughan 2007 and Maynard et al. 2019). For analyses, we calculated the maximum and average percentages of each *Piper* species present in the diet of all three *Carollia* species from this dataset (Table S5).

To investigate if fruit scent composition could be a potential factor explaining differences in *Carollia-Piper* interaction across habitats, we used a chemical dataset of *Piper* ripe fruit VOCs collected at La Selva and published by one of us (Santana et al. 2021). In that study, VOC emission data were obtained from ripe fruits for 21 *Piper* species via headspace adsorption methods and gas chromatography-mass spectrometry (GC-MS). GC-MS peaks were integrated, standards were run to verify peak identities, enantiomers were combined, VOCs matched to ChemStation library compounds, and finally compounds were verified by Kovats Retention Indices. Contaminants and all VOCs present in fewer than five samples were removed from that dataset (see Santana et al. 2021 Supplementary Information).

To compare the fruit scent composition of forest against gap *Piper*, we classified all 21 species in the Santana et al. 2021 dataset into habitat categories, for a total of 13 forest and 8 gap species. We sorted their total VOC emissions per weight for 249 volatile organic compounds for each species, resulting in a list of the most abundant chemicals in each species in order of decreasing concentration. Then, we identified the 15 most common chemical compounds among the chemicals in highest abundance of each of the species (this number was chosen to include most of the *Piper* species in the dataset and avoid null data values, as the number of VOCs among species ranged from 4 to 104 compounds). Additionally, we compiled the total VOC emission and total number of VOCs across the *Piper* species. We used each of these three datasets (abundances of 15 common VOCs, total emission, total number of VOCs; Table S6) in statistical analyses.

### Statistical analyses

We performed all statistical analyses in R v. 4.3.1 (R Core Team 2023). We tested for phylogenetic signal in the data using the time-calibrated, species-level *Piper* phylogeny published in Santana et al. (2021) and the function ‘physignal’ in the package geomorph (Adams et al. 2023). To compare visitation and consumption across *Piper* habitats (open vs. gap; earl-, mid-and late-succession), we used Pearson’s Chi-squared Test for Count Data (Pearson, 1900) in the package stats (R Core Team 2023) and adjusted for 2000 replicates because of our small sample size (Hope 1968).

To linearize the sigmoid distribution of proportions of the diet dataset, we added an arbitrary constant (c = 1) to avoid zero values, logit transformed (y = ln(p/(1-p)) these data (Armitage and Berry 1994) and performed Shapiro-Wilk’s tests (Shapiro and Wilk 1965). These tests indicated that the transformed diet data for *C. castanea, C. sowelli*, and *C. perspicillata* followed normality (W = 0.4216, P = 9.047e-10; W = 0.42227, P = 2.789e-10; W = 0.54719, P = 6.164e-09), which was also the case for the maximum and average percentages of *Piper* in *Carollia* diets (W = 0.51649, P = 2.745e-09; W = 0.49507, P = 1.593e-09). We then performed analyses of variance (ANOVAs [Girden 1992]) to test for differences in the transformed percentages of *Piper* species (forest or gap, and early-, mid-, or late-succession) in *Carollia* diets. To test for differences in fruit scent between forest and gap *Piper*, we used the npmv package (Burchett et al. 2017) for all multivariate analyses, as our data did not follow normality after being tested with a Shapiro-Wilk’s test.

## Results

### Patterns of Carollia-Piper interactions across biomonitoring modalities

The individual methods used to detect frugivores in relation to *Piper* plants had an influence in the type of information that could be retrieved about their interactions, and therefore the conclusions that could be made about habitat patterns. For example, at one end of the spectrum, passive acoustic recording data (in the form of identified echolocation calls) are informative of general bat activity and/or presence of bats near plants, whereas fecal samples directly collected from bats can confirm whether this general bat activity includes fruit consumption that would lead to seed dispersal. Somewhere in between, camera trap video data provides information about plant visitation, and fruit exploratory and procurement behaviors (as *Carollia* do not feed at *Piper* plants directly but take the fruits to a feeding roost first [Wilson and Mittermeier, 2019]).

### (a) Videos

Our camera traps allowed us to document *Carollia* collecting fruit at the forest species *P. cyanophyllum, P. generalense, P. nudifolium,* and *P. sublineatum*, and the gap species *P. colonense, P. reticulatum,* and *P. sancti-felicis* (Table 1, Supplemental Video 1). We observed *Carollia* visitations (flying by, inspecting fruits before leaving) at the forest species *P. generalense, P. nudifolium, P. paulowniifolium, P. sublinateum,* and the gap species *P. reticulatum, P. sancti-felicis,* and *P.* species D. Additionally, we were able to document *Piper* plant visitations and fruit consumptions by insects, birds, and small mammals other than bats (Table 1). Larger animals, such as tapirs, were recorded consuming entire *P. nudifolium* plants as they walked by. Rodents and possums were recorded passing by the cameras or climbing on the plants (F.V. personal observation; Supplemental Video 2). Birds would sometimes perch on the branches without consuming fruits. Based on this range of observations, we classified videos into different behaviors that involved *Piper* fruits: inspecting fruits, removing fruit, and eating fruit. We found bats and birds to most commonly take fruit off of the plants, although some birds ate the fruits while they remained attached to the plant (Supplemental Video 3). Fruit removal/consumption by non-bat frugivores only occurred at gap *Piper*, which were also consumed by birds and insects, whereas targeted collection of fruits by bats only occurred in forest *Piper*. This lends support to our initial hypothesis that frugivore diversity is dependent on *Piper* habitat. We performed a chi-square test of independence on the number of interactions between *Piper* plants and *Carollia* bats identified by camera traps against *Piper* habitat (forest and gap) and failed to reject the null hypothesis (with 2000 replicates; P = 0.2084). However, a chi-square test using a finer *Piper* habitat classification (early-, mid-, late-succession) on the same data resulted in significant differences (with 2000 replicates: X^2^=8, df=NA, p-value=0.01949).

The camera trap data produced additional insight into the general activity patterns of *Carollia* visiting and consuming *Piper* over the course of the night and throughout the year. As seen in Fig. 4, general fruit acquisition and visitation activity by bats is continuous from dusk throughout the night until dawn, with a peak earlier in the night. We also documented more frequent visitations to gap *Piper* earlier in the night (Fig. 4) and observed a difference in the number of *Piper* species in which bat activity was recorded throughout the night (Fig. 5); bats visit a greater number of gap *Piper* species early in the night, and fewer species later on. This pattern was not seen at forest *Piper* plants, where bats visited a different forest *Piper* species every hour or so, but not more than one. Throughout the year (excluding August, September, and October, as we did not record field data during this time), we observed bats taking fruit and visiting both forest and gap *Piper*.

**Figure 5.**
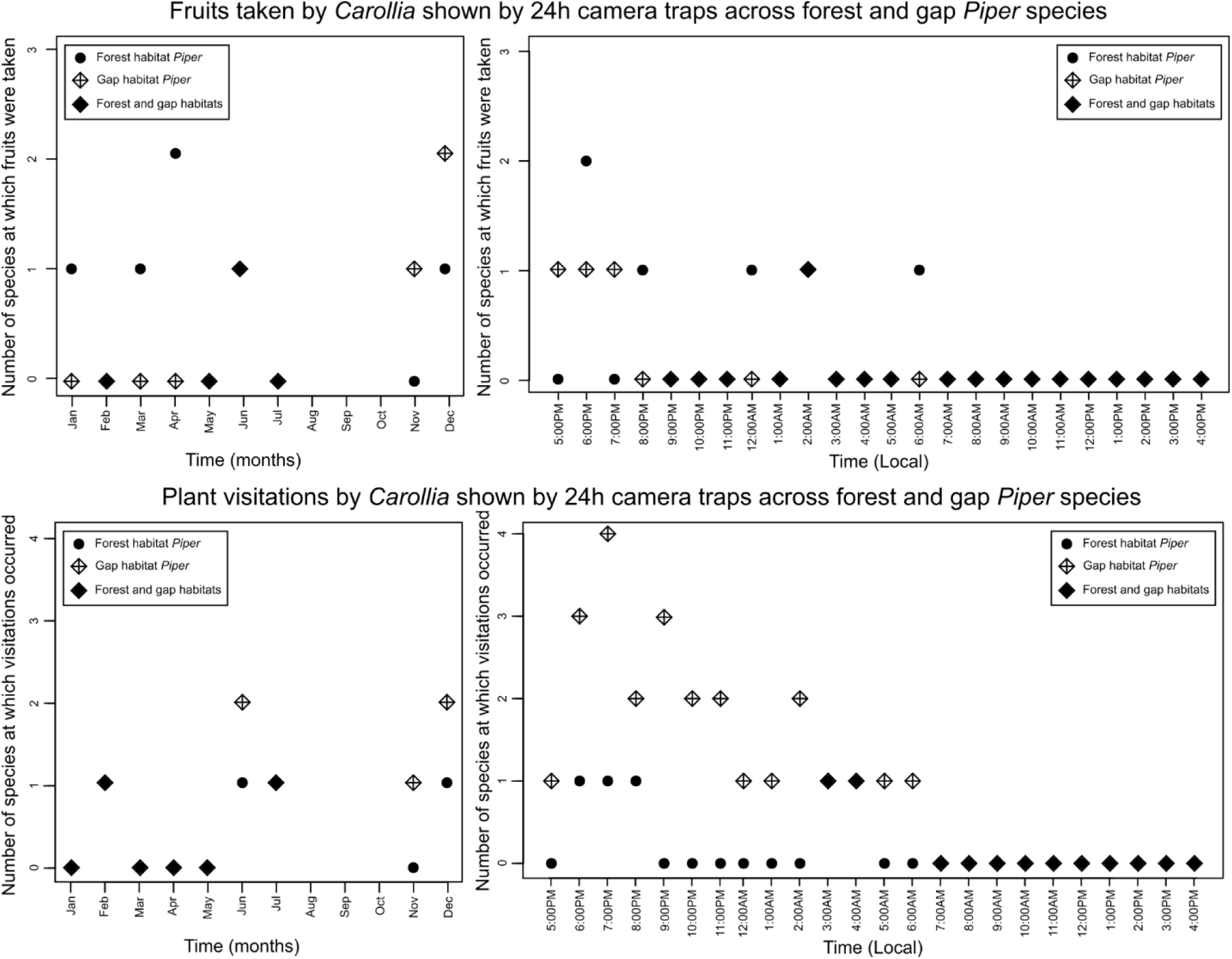
Temporal patterns of bat activity across forest and gap *Piper* species. Top: Total number of species at which instances of bats taking fruits occurred as shown by 24h camera traps across forest and gap *Piper* over the course of the year and throughout the day. Data are shown starting from sunset (5:00 PM local time), when bat foraging begins. Bottom: Total number of species at which *Carollia* visitations to plants were recorded as shown by 24h camera traps across the same temporal scales. Data are the visitations and fruit acquisitions recorded as a binary at each *Piper* species where camera data revealed bat activity, indicating the number of *Piper* species (classified by habitat) where activity was recorded at a certain time or during a month.

### (b) Acoustics

Acoustic monitoring allowed us to document the presence of bats at the forest *Piper* species *P. generalense, P. nudifolium,* and *P. paulowniifolium*, and the gap species *P. reticulatum*. Collection of *Piper* fruits by bats could not be identified purely by this method. However, we identified search-phase echolocation calls with harmonics, which indicate *Carollia* bats flying by, if not possibly visiting *Piper* plants to inspect fruits. We performed chi-square tests of independence on the acoustic visitation results with *Piper* habitat (open, forest, or early-, mid-, or late-succession, respectively) as predictor variables; the relationship between these two variables was not significant in both cases (with 2000 replicates each: P = 0.2239; P = 0.9999).

While we detected *Carollia* calls in our dataset, our results were dominated by insectivorous bats in the vicinity of *Piper* plants. These calls were identified by the presence of FM-QCF (frequency-modulated, quasi-constant frequency), FM-CF (frequency-modulated, constant frequency), FM-CF-FM (frequency-modulated, constant frequency, frequency-modulated) calls and feeding/terminal buzzes (see Fig. 3, Table S4). Additionally, we found numerous calls of the common vampire bat (*Desmodus rotundus*) in our data due to the presence of a large roost near one of the sites (S.E.S. personal observation). The key differences between the calls of *Carollia* and insectivorous bats’ are that *Carollia*’s calls exhibit higher frequencies and purely frequency-modulated components. *Carollia* and *D. rotundus* calls were distinguished by their higher and lower frequency bandwidths, respectively (Collen 2012).

### (c) Diet

We used ANOVAs to compare the maximum and average percentages of *Piper* in *Carollia* diets against the habitat classifications as predictor variables. These analyses resulted in statistically significant differences in consumption of *Piper* species when these were categorized by their habitat (two-and three-category schemes; p < 0.001, see Tables 2a and 2b respectively). The results further provide evidence that all three *Carollia* species consume significantly more gap (early-succession) *Piper* than forest (mid-, late-succession) *Piper*. Nonparametric inference for the comparison of multivariate data samples (Burchett et al. 2017) testing the aforementioned variables indicated that there is a 95% probability that a randomly chosen *Carollia* exhibits a larger percentage of gap than forest *Piper* in their diet.

**Table 2.** One-way analyses of variance (ANOVAs) testing the percent of *Piper* species in the diets of the three *Carollia* species against habitat as a predictor variable (following logit transformation and testing for normality). (*** P < 0.01).

| A. Predictor variable: Habitat (forest, gap) |  |  |  |  |  |
| --- | --- | --- | --- | --- | --- |
|  | df | Sum sq | Mean sq | F-value | P |
| *C. castanea* | 1 | 0.04763 | 0.04763 | 14.08 | 0.000725*** |
| *C. sowelli* | 1 | 0.06412 | 0.06412 | 13.63 | 0.000852*** |
| *C. perspicillata* | 1 | 0.05261 | 0.05261 | 23.76 | 3.08e-05*** |
| Max. % | 1 | 0.09305 | 0.09305 | 19.63 | 0.000109*** |
| Average % | 1 | 0.05363 | 0.05363 | 18.38 | 0.000164*** |
| B. Predictor variable: Succession habitat (early, mid, late-succession) |  |  |  |  |  |
|  | df | Sum sq | Mean sq | F-value | P |
| *C. castanea* | 2 | 0.05356 | 0.026782 | 8.119 | 0.00152** |
| *C. sowelli* | 2 | 0.06858 | 0.03429 | 7.279 | 0.00265** |
| *C. perspicillata* | 2 | 0.05534 | 0.27671 | 12.59 | 0.000107*** |
| Max. % | 2 | 0.1037 | 0.05183 | 11.41 | 0.000207*** |
| Average % | 2 | 0.05772 | 0.028861 | 10.03 | 0.000463*** |

### *Fruit scent composition as a medium for interpreting* Carollia-Piper *habitat patterns*

The variation in the fruit scent VOC data used in our analyses was not highly impacted by the evolutionary relationships between *Piper* species; there was no significant phylogenetic signal for almost all of the first 15 most common VOCs, with the exception of the most common VOC across the *Piper* species in the dataset, alpha-caryophyllene, which approached significance (Pagel’s K = 0.83, P = 0.053). Results from non-parametric multivariate tests indicated significant differences in the chemical composition of forest versus gap *Piper* (test statistic = 3.126, df1 = 6.004, df2 = 106.1721, P = 0.007, permutation test P = 0.006). Pairwise comparisons using a Wilcoxon rank sum test (Wilcoxon 1945) with continuity correction (p-value adjustment method: Benjamini and Hochberg [Benjamini and Hochberg 1995]) yielded significant differences between gap versus forest *Piper* (Fig. 6) for the VOCs beta-pinene (P = 0.0052), 2-dodecene (P = 0.046), beta-elemene (P = 0.011), 3-methyl-2-undecene (P = 0.038), 3- methyl-3-undecene (P = 0.038), and decanal (P = 0.014). We did not find differences in the total emission and number of VOCs among *Piper* species classified by habitat (P = 0.244; P = 0.153) or succession stage (P = 0.663; P = 0.074).

**Figure 6.**
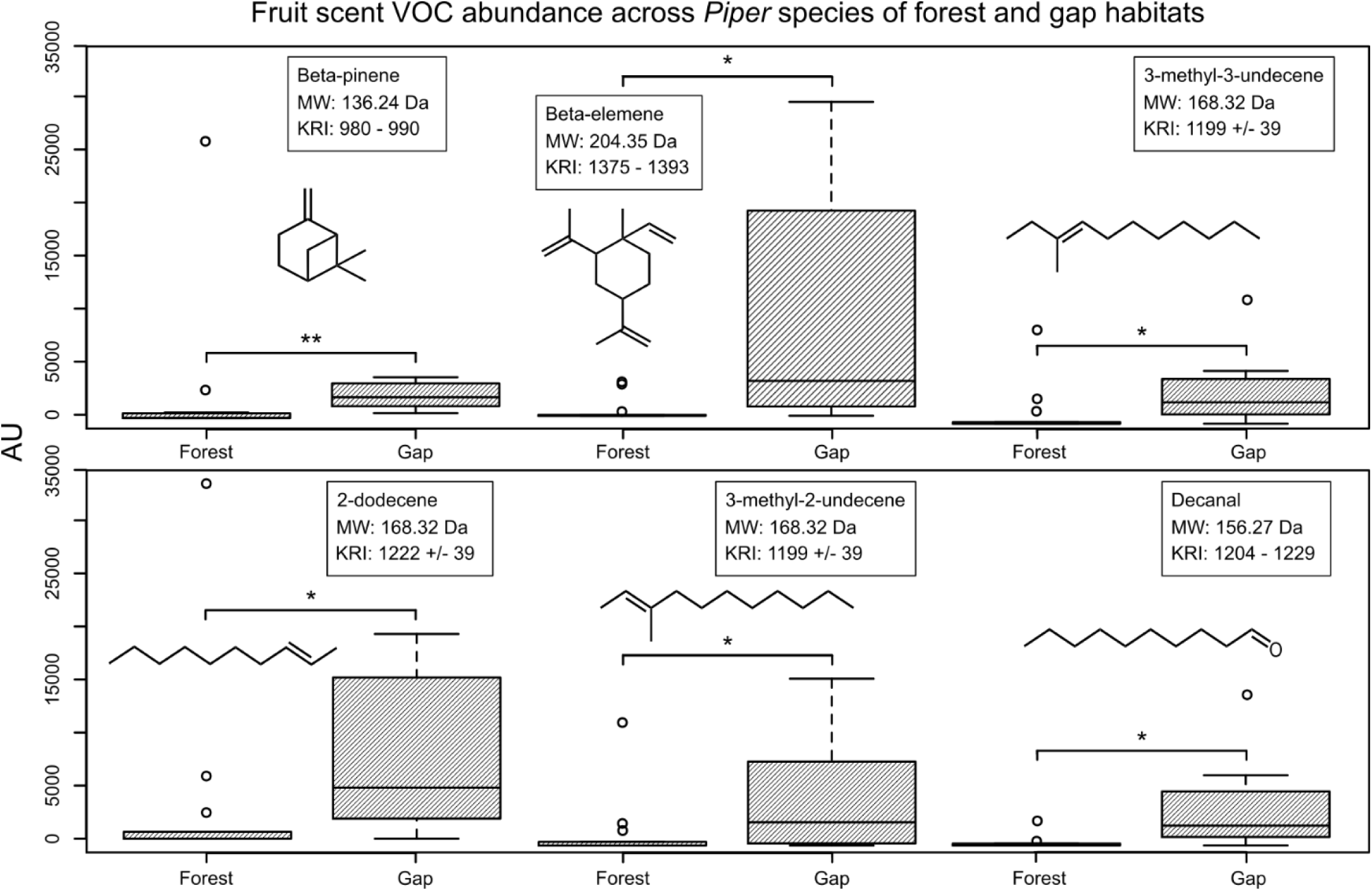
Six chemicals that are significantly different between gap and forest *Piper* fruit scent compositions (beta-pinene, beta-elemene, 3-methyl-3-undecene, 2-dodecene, 3-methyl-2- undecene), showing the difference in VOC emissions per weight between forest and gap species (*: P < 0.05, **: P < 0.01). For reference, chemical structures, molecular weights (MW), and Kovats Retention Indices (KRI) are included for each chemical. KRIs for 2-dodecene, 3-methyl-2-undecene, and 3-methyl-3-undecene were referenced from ChemSpider. KRIs for beta-pinene, beta-elemene, and decanal were reported as ranges for column type DB-5 (matched with the methods reported in Santana et al. 2021) collected from the literature sources compiled in The Pherobase: Database of Pheromones and Semiochemicals.

## Discussion

### Different insights are gained from acoustic monitoring, video data, and diet studies when characterizing plant-bat interactions

Our study aimed to document frugivore visitation patterns across putative *Piper* ecotypes in Costa Rica to gain more insight into the *Carollia-Piper* mutualism, and henceforth allow for a better understanding of biodiversity and behavioral ecology in the Neotropics. We integrated three biomonitoring modalities to devise the best approach to characterize these fruit-bat interactions and interpret the relationship between Neotropical *Piper* and short-tailed fruit bats.

Our analysis of nightly acoustic recordings allowed us to identify frugivorous, insectivorous, and sanguinivorous bat activity at or near *Piper* plants. In this case, bat activity should not be confused with abundance (Fraser et al. 2020) or any type of specific behavior, however, so these results only indicate the presence of *Carollia* bats at or near a *Piper* plant. Regardless, the number of *Carollia* visitations found by our acoustic analyses was not sufficiently large for determining differences in bat presence at one *Piper* habitat versus another. Since *Carollia* have low-intensity calls characteristic of phyllostomids, our acoustic recorders were turned up to high gain, leading to a high level of background noise in the recordings. After filtering, we obtained clear evidence of general bat activity at most of the *Piper* species and plants (Fig. 3, Table S4), ruling out the possibility of our recorders failing to detect ultrasonic signals. Given that *Carollia* are some of the most abundant bats at La Selva, technical challenges probably affected our ability to successfully record clear *Carollia* search-phase echolocation calls. Additionally, previous behavioral studies have established that *C. castanea* and *C. perspicillata* initially find ripe fruit primarily using olfaction and only rely more heavily on echolocation when olfactory cues are insufficient (Thies et al. 1998; Leiser-Miller et al. 2021), providing further explanation for the difficulty in capturing *Carollia* visitations to *Piper* through acoustic methods.

The scarcity of *Carollia* echolocation calls found in the passive acoustic monitoring field data compared to the abundance of detected insectivorous bat activity directs us to look into additional physical factors that could have affected these results. Bat species that emit broad bandwidth calls like *Carollia* are more successful in detecting details and therefore finding food items in cluttered environments, but the high frequencies that dominate these FM calls undergo significant atmospheric attenuation, so these calls are only effective and detectable by acoustic devices over a short range (Griffin 1971; Lawrence and Simmons 1982; Neuweiler 1990). *Carollia* calls are low intensity and high frequency, and attenuate very quickly in warm, humid environments (e.g., 45–90 kHz attenuates at 1.4–4 dB/m at 25°C and 80% humidity [Jakobsen et al. 2013; Leiser-Miller et al. 2021]). To provide further support for this phenomenon, we did not detect any calls above 70-80kHz in our recordings (S.Sil personal observation). Bats also tend to increase call frequency and pulse rates and decrease the duration of their calls to avoid forward masking, which is essential in cluttered environments and dense vegetation (Schnitzler and Kalko 2001). In summary, distance, clutter, and noise all affect the quality of acoustic recordings in the field (Fraser et al. 2020); it is up to the discretion of the scientists wishing to monitor bat activity in their chosen field site to decide at which taxonomic level they are comfortable identifying (based on the available call libraries and focal data), how they will filter the noise from their recordings, and how to account for phenomena such as atmospheric attenuation in their results.

Camera traps are considered a minimally invasive biosurveying method when compared to those requiring capture or human presence, such as mist-netting or visual counts (Sollmann et al. 2013; Krivek et al. 2021). However, bats, due to their small size, nocturnal habits, and ability to fly quickly (Krivek et al. 2021) have been a challenge to monitor with traditional methods like photo-based camera traps. In our study, and thanks to technical advances in detection speed and infrared recording, video-based camera traps provided a better understanding of interactions between *Carollia* and *Piper*: by examining video recordings, we were able to directly see *Carollia* taking *Piper* fruit and *Carollia* inspecting fruits for some time before grabbing one (or not) and flying away. We were also able to see a variety of other animals interacting with *Piper* plants, providing a unique “plant perspective” of the interaction. Due to this functionality of the camera traps, we were able to find support for the hypothesis that forest *Piper* species depend on *Carollia* as their main seed dispersers during their shorter fruiting periods, whereas gap species exhibit a broader range of dispersers, most of which are still bats (see Table 1). It’s important to note, however, that we found statistical support for these differences only when *Piper* were classified into finer habitat categories (early-, mid-, late-succession). This discrepancy may have resulted from the fact that many mid-succession *Piper* species do not fall neatly in a forest vs. gap habitat categorization, but rather in between. Combing through thousands of hours’ worth of videos is still an arduous task (automated methods proved inefficient due to wind-or rain-caused plant movement and other environmental factors) and may not provide the exact information needed to characterize some bat-plant interactions. For example, in our system, some interactions were never detected despite the *Piper* species being reported as one significant component of *Carollia’s* diet (e.g., gap species *P. umbricola;* average 8.51% of the diet of three *Carollia* species [Santana et al. 2021]). While the reasons why the bats never visited the selected plants of these species during our study are not clear, this issue could be solved by increasing the sample size and duration of the study.

Examining which *Piper* species are present in fecal samples collected in the field is the most direct way to measure consumption of *Piper* by *Carollia*. The *Carollia-Piper* mutualism hinges on the regular ingestion of *Piper* fruit (including whole seeds) by *Carollia* in tandem with variation in the fruiting peaks roughly characteristic to gap and forest *Piper*; forest *Piper* species rely on *Carollia* for seed dispersal during their short fruiting period, and gap *Piper* provide nutrients year-round for *Carollia* (e.g., *P. sancti-felicis* [Leiser-Miller et al. 2020; Santana et al. 2021; Maynard et al. 2019]). The results of diet data analyses were consistent with those from video data in uncovering significant differences in *Carollia* interactions across *Pipers* of different habitats, even at the coarser levels of habitat classification. This suggests that the more intensive and direct approach of collecting fecal samples brings us closer to discovering habitat effects in this *Carollia-Piper* mutualism. However, it is important to consider that analyses of bat fecal samples for dietary identification at this scale is labor-intensive both in the field and the lab, and requires seed reference libraries collected over months or years.

By capitalizing on annual diet data based on fecal samples collected from hundreds of bats, we were able to describe that *Carollia* (*C. sowelli, C. perspicillata, C. castanea*) consume a greater percentage of gap *Piper* species than forest *Piper* species. As proposed by Thies and Kalko (2004), phenology can provide an explanation for this phenomenon; forest *Piper* species fruit for a relatively shorter time period than gap *Piper*, and therefore gap *Piper* consumption will be higher on average when considered throughout the year. For this reason, cross-checking with visitation or consumption data on smaller temporal scales (e.g., during the same night across species and habitats), as can be done by camera trap data analysis could help gain more insights regarding the finer scale dynamics of the *Carollia-Piper* mutualism. At present, our camera trap data are not sufficient to do so, since there were few bat visits/consumption events that coincided between forest and gap *Piper* in the same night; future studies could use the methods presented here across a greater number of plants in selected *Piper* species to look at these patterns.

Generally, bat biologists collect bats and their fecal samples early in the night when bats are known to be foraging most actively. Our camera results provide evidence that *Carollia* could be visiting and consuming a greater number of *Piper* species –particularly gap species– earlier in the night, followed by decreased activity and a switch to consumption of forest *Piper* later in the night (Figs. 4 and 5). These results expand on the findings of Heithaus and Fleming, who noted *C. perspicillata* activity throughout the night (Heithaus and Fleming 1978) and highlight the need to extend *Carollia* behavioral and diet surveys later into the night to capture its full diet diversity. Overall, our results indicate that analyzing diet datasets synergizes well with real-time monitoring of plants and bat activity, however; by combining acoustic and video methods, we gained insight into plant-bat interactions across different levels: bat presence around plants, visitations to plants, inspection of plants, and acquisition and consumption of fruits.

### *Differences in* Piper *scent volatiles may influence* Carollia-Piper *interactions across habitats*

Phyllostomid frugivores integrate across sensory modalities to locate and acquire ripe fruit, using vision to detect fruit color, olfaction to detect fruit scent volatiles, and echolocation to collect information on the location and shape of fruits (Schwab and Pettigrew 2005; Hodgkison et al. 2013; Kalko and Condon 1998; Von Helversen and Von Helversen 1999; Leiser-Miller et al. 2021; Santana et al. 2021). Previous research suggests that frugivorous phyllostomids such as *Carollia* modulate their behavior with respect to olfaction and echolocation to maximize the effectiveness of one sensory cue versus the other when appropriate (Thies and Kalko 2004; Leiser -Miller et al. 2021). Further, previous studies have shown *Carollia* mainly rely on scent cues (volatile organic compounds found in fruit scent composition) for selecting ripe *Piper* fruits (Thies et al. 1998; Leiser-Miller et al. 2020; Santana et al. 2021). Therefore, in order to mechanistically understand *Carollia-Piper* interactions, it is informative to examine the differences between forest and gap *Piper* in terms of their fruit scent chemical composition. In particular, the extent of ripeness and type and strength of the chemical cues could be key to affecting bat visitations and behaviors, as a ripe fruit with a strong signal could be located and seized very quickly, whereas a fruit still ripening or with weak signals may end up uneaten even after a long period of inspection.

We find evidence for differences in the chemical composition of fruit scent between forest and gap *Piper* species (Fig. 6); via olfactory preferences, these distinguishing chemicals could potentially underlie differences in *Carollia* visitation and consumption to and of *Piper* across habitats. We identified six VOCs among the most common chemicals found in the scent bouquet of 21 *Piper* species to be significantly more abundant in gap *Piper* compared to forest *Piper*. Two are terpenes (one monoterpene, one sesquiterpene), three are long hydrocarbon (C_n_ = 10, C_n_ = 11) chain alkenes, and one is a long hydrocarbon chain (C_n_ = 10) aldehyde. Studies have shown that mammals, including bats, have a higher olfactory performance (sensitivity) when tested on compounds containing longer carbon chains (Laska, Seibt and Weber 2000); hence, even low concentrations of these compounds in the scent bouquets of *Piper* are likely to attract bats to the fruit (Borges et al. 2008). Our results (long hydrocarbon chain compounds found to be more abundant in the emissions of gap *Piper* fruits compared to forest), along with these studies, provide further evidence that *Piper* fruit scents may influence *Carollia* visitations across habitats. Additionally, Santana et al. found that highly consumed *Piper* species, which are included in our dataset, are phylogenetically scattered and characterized by scents rich in terpenoids, similar to other bat-dispersed fruits (which contain high abundances of monoterpenes [Parolin et al. 2019; Hodgkison et al. 2013; Santana et al. 2021]). Our results, which include beta-pinene and beta-elemene (terpenoids characteristic to gap *Piper* species), support these findings. Importantly, the fruit scent of a *Piper* species highly consumed by *Carollia* (*P. sancti-felicis*) is also unique in containing 2-heptanol, an aliphatic alcohol preferred by *Carollia* in behavioral experiments (Leiser-Miller et al. 2020; Santana et al. 2021). Thus, particular notes in the fruit scent bouquet may also play a role in *Piper* preferences. Further behavioral experiments are necessary to determine if and which fruit scent chemicals contribute to driving differences in *Piper* consumption across habitats; the aforementioned terpenoids and hydrocarbon chain compounds we identified through our analyses are good candidates for this future work.

## Conclusion

Our results integrating acoustic, camera trap, and diet data lend support to the hypothesis that forest and gap *Piper* differ in their diversity of interacting frugivores, with forest *Piper* exhibiting a tight relationship with bats, and gap *Piper* interacting with a wider spectrum of frugivores. We found that, for the three *Carollia* species present at our study site (*C. sowelli, C. perspicillata, C. castanea*), gap *Piper* was consumed significantly more than forest *Piper*, but visitations and fruit acquisition by bats occurred across both forest and gap *Piper* throughout the year, and forest *Piper* were only visited by *Carollia*. We observed *Carollia* visitations to gap and forest *Piper* throughout the night, but also found evidence for a foraging activity peak closer to dusk characterized by a greater variety of gap *Piper* species visited or collected by bats. By incorporating fruit scent chemical data into our analyses of *Piper* ecotypes, we not only find support for the hypothesis that scent signals might drive differential foraging by *Carollia* on *Piper* fruits, but we identify specific compounds (terpenoids, hydrocarbon chain derivatives) that may influence *Carollia* visitations across forest and gap habitats. Our study highlights the benefit of integrating multiple biomonitoring methods and datasets to characterize plant-animal interactions.

## Authors’ contributions

S.S. formulated and carried out analysis of the acoustic recording dataset and performed statistical analyses with input from S.E.S. F.V. performed camera trap video analysis. G.C. contributed to study design and fieldwork. S.E.S. designed the study and conducted all field data collection. S.S. and S.E.S. interpreted the data and wrote the first draft of the manuscript. All authors revised the manuscript and gave final approval for publication.

## Competing interests

We have no competing interests.

## Acknowledgements

Orlando Vargas Ramirez, Danilo Brenes Madrigal, Bernal Matarrita Carranza, Enrique Castro Fonseca, and staff at La Selva Biological Station provided invaluable support during fieldwork, particularly facing the challenges of international fieldwork increased by the COVID-19 pandemic. Anusha Aggarwal, Betsaida Rodriguez, Dr. Iroro Tanshi, and Dr. Wu-Jung Lee contributed to the conception of our acoustic analysis method, and Dr. Priya Balasubramanian supported the development of our analytical algorithm with her expertise in signal processing.

## Funding sources

This work was supported by the Mary Gates Endowment for Students [Mary Gates Research Scholarship to S.S.]; the Endowment for the Thomas Sedlock Icon Scholarships [Thomas Sedlock Icon Scholarship to S.S.]; and the Fulbright Scholars Program [Costa Rica fellowship to S.E.S.].

## Notes

### Competing Interest Statement

The authors have declared no competing interest.

