## Supplemental Information for "Effects of habitat and fruit scent on the interactions between short-tailed fruit bats and *Piper* plants"

Supplemental materials

Code for MATLAB acoustic analysis algorithm available upon request.

Supplementary Video 1: Bat flying past *Piper* plant and acquiring fruit.

Supplementary Video 2: Mouse climbing on a *Piper generalense* plant

Supplementary Video 3: Passerini's tanager consuming *Piper sancti-felicis*.

Supplemental tables

**S1.** Recorder locations, plant site locations, # of ripe fruits, starting and end times/dates

**S2.** Focal call parameters from search-phase echolocation calls of three *Carollia* species

**S3.** Number of acoustic files identified as containing bat calls via acoustic algorithm

**S4.** Number of each type of bat echolocation call identified within the acoustic dataset

**S5.** Diet dataset from the literature (Santana et al., 2021; Lopez & Vaughan, 2007; Maynard et al., 2019) with *Piper* habitat classifications

**S6.** Chemical dataset sorted by concentrations per *Piper* species of the 15 most abundant chemicals among all *Piper* species published in Santana et al. 2021

**Table 1.** Information on the *Piper* plants studied in the field, their locations, the dates and number of days recorded (audiomoths and camera traps).

| Species | Habitat | Classification | Site | # ripe fruits | Duration (days) | Start date | End date | LAT (N) | LONG (W) |
|---|---|---|---|---|---|---|---|---|---|
| *P. auritum* | gap | early-suc | 1 | 2 | 36 | 6/5/2021 | 7/11/2021 | 10.43146 | -84.00369 |
| *P. colonense* | gap | mid-suc | 1 | 3 | 12 | 12/1/2019 | 12/13/2019 | 10.4308 | -84.00645 |
| | | | 2 | 4 | 11 | 6/4/2021 | 6/15/2021 | 10.43089 | -84.00334 |
| | | | 3 | 3 | | 6/15/2021 | | 10.43015 | -84.00971 |
| *P. cyanophyllum* | forest | mid-suc | 1 | 1 | 139 | 1/26/2020 | 6/13/2020 | 10.4311 | -84.00666 |
| *P. generalense* | forest | mid-suc | 1 | 3 | 84 | 11/16/2019 | 2/8/2020 | 10.42501 | -84.0016 |
| | | | 2 | 2 | 84 | 11/16/2019 | 2/8/2020 | 10.42502 | -84.00171 |
| | | | 3 | 5 | 70 | 11/16/2019 | 1/25/2020 | 10.42502 | -84.0017 |
| | | | 4 | 2 | 14 | 11/16/2019 | 11/30/2019 | 10.42483 | -84.00164 |
| | | | 5 | 2 | 70 | 11/16/2019 | 1/25/2020 | 10.43071 | -84.00674 |
| | | | 6 | 3 | 96 | 11/17/2019 | 2/21/2020 | 10.4335 | -84.01216 |
| | | | 7 | 7 | 211 | 11/17/2019 | 6/15/2020 | 10.43377 | -84.01219 |

| | | | | | | | | | |
|---|---|---|---|---|---|---|---|---|---|
| | | | 8 | 14 | 135 | 1/26/2020 | 6/9/2020 | 10.43733 | -84.00581 |
| *P. multiplinervium* | gap | early-suc | 1 | 5 | 69 | 11/16/2019 | 1/24/2020 | 10.4306 | -84.00658 |
| *P. nudifolium* | forest | mid-suc | 1 | 3 | 140 | 1/27/2020 | 6/15/2020 | 10.43543 | -84.03122 |
| *P. paulowniifolium* | forest | mid-suc | 1 | 7 | 35 | 6/6/2021 | 7/11/2021 | 10.4328 | -84.00667 |
| | | | 2 | 11 | 35 | 6/6/2021 | 7/11/2021 | 10.43854 | -84.01363 |
| | | | 3 | many, count in photo | 25 | 6/16/2021 | 7/11/2021 | 10.43749 | -84.01147 |
| *P. reticulatum* | forest | early-suc | 1 | 18 | 44 | 11/17/2019 | 12/31/2019 | 10.43821 | -84.01594 |
| | | | 2 | 16 | 16 | 11/27/2019 | 12/13/2019 | 10.43045 | -84.0059 |
| | | | 3 | 14 | 12 | 6/3/2021 | 6/15/2021 | 10.43699 | -84.01009 |
| | | | 4 | 3 | 12 | 6/3/2021 | 6/15/2021 | 10.43642 | -84.00952 |
| | | | 5 | 6 | 6 | 6/4/2021 | 6/10/2021 | 10.43295 | -84.00443 |
| | | | 6 | 4 | 9 | 6/6/2021 | 6/15/2021 | 10.43446 | -84.00821 |
| | | | 7 | 3 | 13 | 6/10/2021 | 6/23/2021 | 10.43053 | -84.00675 |

| | | | | | | | | | |
|---|---|---|---|---|---|---|---|---|---|
| | | | 8 | 5 | 8 | 6/10/2021 | 6/18/2021 | 10.43015 | -84.00971 |
| | | | 9 | 11 | 25 | 6/16/2021 | 7/11/2021 | 10.43069 | -84.00661 |
| | | | 10 | 9 | 18 | 6/23/2021 | 7/11/2021 | 10.42969 | -84.00836 |
| *P. sanctifelicis* | gap | mid-suc | 1 | 2 | 3 | 11/16/2019 | 11/19/2019 | 10.43115 | -84.00694 |
| | | | 2 | 2 | 5 | 11/19/2019 | 11/24/2019 | 10.431 | -84.00646 |
| | | | 3 | 2 | 3 | 11/24/2019 | 11/27/2019 | 10.43076 | -84.00656 |
| | | | 4 | 1 | 1 | 12/1/2019 | 12/2/2019 | 10.43043 | -84.00608 |
| | | | 5 | 1–3 | 8 | 12/5/2019 | 12/13/2019 | 10.43075 | -84.00682 |
| | | | 6 | 3 | 9 | 1/27/2020 | 2/5/2020 | 10.43005 | -84.01115 |
| | | | 7 | 3 | 17 | 2/5/2020 | 2/22/2020 | 10.43084 | -84.00675 |
| | | | 8 | 7 | 5 | 2/12/2020 | 2/17/2020 | 10.43148 | -84.00584 |
| | | | 9 | 1 | 4 | 6/6/2021 | 6/10/2021 | 10.43136 | -84.00648 |
| | | | 10 | loaded! See photo | 23 | 6/18/2021 | 7/11/2021 | 10.42989 | -84.00835 |
| | | | 11 | 6 | 13 | 6/28/2021 | 7/11/2021 | 10.43322 | -84.00737 |
| *P. species D* | gap | mid-suc | 1 | 3, many more (count in photo) | 31 | 1/26/2020 | 2/26/2020 | 10.43052 | -84.00613 |

| | | | | | | | | | |
|---|---|---|---|---|---|---|---|---|---|
| | | | 2 | 3 | 16 | 2/10/2020 | 2/26/2020 | 10.43132 | -84.0061 |
| | | | 3 | 4 | 12 | 6/15/2021 | 6/27/2021 | 10.43104 | -84.00624 |
| *P. sublinateum* | forest | mid-suc | 1 | 2 | 40 | 2/23/2020 | 4/3/2020 | 10.43493 | -84.00412 |
| | | | 1.2 | 2 | 1 | 6/26/2021 | 6/27/2021 | 10.43493 | -84.00412 |
| *P. umbricola* | gap | early-suc | 1 | many, count in photo | 30 | 1/26/2020 | 2/25/2020 | 10.43183 | -84.006 |
| | | | 2 | 3 | 8 | 2/18/2020 | 2/26/2020 | 10.43153 | -84.00594 |

**Table 2.** Parameters of search-phase echolocation call focal data of *C. castanea*, *C. perspicillata*, and *C. sowelli* calculated using RavenPro v. 1.6.2's spectrogram display and analysis capabilities (512 FFT Hanning window, 95% overlap; see Methods).

| **Species** | **N** | | **Delta Time (ms)** | **Dur 90% (ms)** | **Low Freq (kHz)** | **High Freq (kHz)** | **Freq 95% (kHz)** | **Peak Freq (kHz)** | **Delta Freq (kHz)** | **BW 90% (kHz)** |
|---|---|---|---|---|---|---|---|---|---|---|
| *C. castanea* | 255 | Average | 1.18 | 0.615 | 66.3 | 95 | 91.7 | 80.2 | 28.7 | 20.4 |
| | | Standard deviation | 0.295 | 0.286 | 4.94 | 6.34 | 6.2 | 7.87 | 7.78 | 6.35 |
| | | 66% | 0.887 - 1.48 | 0.328 - 0.901 | 61.4 - 71.2 | 88.7 - 101.3 | 85.5 - 97.9 | 72.4 - 88.1 | 20.9 - 36.5 | 14.1- 26.8 |
| | | Min - max | 0.5 - 2.1 | 0.0 - 1.4 | 58.5 - 85.8 | 79.9 - 115.0 | 78.4- 112.0 | 65.9 - 102.0 | 12.0 - 45.6 | 8.79 - 36.6 |
| *C. perspicillata* | 28 | Average | 1.06 | 0.468 | 52.2 | 77.6 | 71.1 | 66 | 25.3 | 13.5 |
| | | Standard deviation | 0.211 | 0.261 | 4.61 | 3.12 | 3.08 | 3.41 | 4.68 | 4.03 |
| | | 66% | 0.849 - 1.27 | 0.207 - 0.729 | 47.6 - 56.8 | 74.5 - 80.7 | 68.0 - 74.1 | 62.6 - 69.4 | 20.6 - 30.0 | 9.48 - 17.5 |
| | | min - max | 0.5 - 1.5 | 0 - 1.0 | 45.1 - 59.6 | 71.5 - 84.8 | 63.5 - 75.4 | 57.6 - 75.2 | 16.7 - 34.7 | 7.81 - 23.4 |
| *C. sowelli* | 69 | Average | 1.28 | 0.386 | 59.7 | 79.2 | 76.4 | 70.9 | 19.5 | 12 |
| | | Standard deviation | 0.33 | 0.371 | 7.08 | 4.43 | 4.42 | 5.02 | 5.81 | 3.22 |

| | | | | | | | | | |
|---|---|---|---|---|---|---|---|---|---|
| | | 66% | 0.952 - 1.61 | 0.015 - 0.756 | 52.6 - 66.8 | 74.8 - 83.6 | 72.0 - 80.8 | 65.9 - 75.9 | 13.7 - 25.3 | 8.78 - 15.2 |
| | | min - max | 0.6 - 2.2 | 0 - 1.4 | 45.0 - 76.8 | 71.0 - 88.6 | 66.7 - 85.7 | 56.4 - 81.3 | 9.50 - 29.8 | 6.59 - 18.3 |

**Table 3.** Number of files identified as likely containing bat calls after filtering by MATLAB signal processing algorithm with more sensitive parameters (Run 1, low pass 100kHz, high pass 20kHz, peak frequency 40kHz) and with more specific parameters (Run 2, low pass 110kHz, high pass 30kHz, peak frequency 50kHz).

| **Species** | **Habitat** | **Site** | **Total files** | **Post Run 1** | **% reduction** | **Post Run 2** | **% reduction** | **N files (Run 1 + Run 2)** |
|---|---|---|---|---|---|---|---|---|
| *P. auritum* | Gap | 1 | 6120 | 466 | 92.39 | | | 466 |
| *P. colonense* | Forest | 1 | 1626 | 54 | 96.67 | 346 | 96.68 | 400 |
| | | 2 | 1440 | 270 | 81.25 | 498 | 65.42 | 768 |
| | | 3 | 9090 | 201 | 97.79 | 923 | 89.85 | 1124 |
| *P. cyanophyllum* | Forest | 1 | 6307 | 123 | 98.05 | 87 | 98.62 | 210 |
| *P. generalense* | Forest | 1 | 6321 | 124 | 98.03 | 171 | 97.29 | 295 |
| | | 2 | 6327 | 126 | 98 | 170 | 97.31 | 296 |
| | | 3 | 6327 | 125 | 98.02 | 172 | 97.28 | 297 |
| | | 4 | 4321 | 87 | 97.99 | 142 | 96.71 | 229 |
| | | 5 | 7099 | 240 | 96.62 | 504 | 92.9 | 744 |
| | | 6 | 6314 | 134 | 97.88 | 222 | 96.48 | 356 |

| | | | | | | | | |
|---|---|---|---|---|---|---|---|---|
| | | 7 | 6314 | 479 | 92.41 | 222 | 96.48 | 701 |
| | | 8 | 6221 | 123 | 98.02 | 87 | 98.6 | 210 |
| *P. multiplinarium* | Gap | 1 | 9360 | 343 | 96.33 | 931 | 90.05 | 1274 |
| *P. nudifolium* | Forest | 1 | 4101 | 90 | 97.81 | 77 | 98.12 | 167 |
| *P. paulowniifolium* | Forest | 1 | 10800 | 470 | 95.64 | 1767 | 83.64 | 2237 |
| | | 2 | 10800 | 1987 | 81.6 | 1767 | 83.64 | 3754 |
| | | 3 | 9000 | 164 | 98.18 | 936 | 89.6 | 1100 |
| *P. reticulatum* | Forest | 1 | 5040 | 89 | 98.23 | 150 | 97.02 | 239 |
| | | 2 | 2362 | 42 | 98.22 | 35 | 98.52 | 77 |
| | | 3 | 1440 | 270 | 81.25 | 706 | 50.97 | 976 |
| | | 4 | 1440 | 150 | 89.58 | 706 | 50.97 | 856 |
| | | 6 | 1440 | 38 | 97.36 | 706 | 50.97 | 744 |
| | | 7 | 4318 | 466 | 89.21 | 803 | 50.97 | 1269 |
| | | 8 | 2470 | 0 | 100 | 789 | 68.06 | 789 |
| | | 9 | 9000 | 164 | 98.18 | 933 | 89.63 | 1097 |
| | | 10 | 6211 | 4 | 99.93 | 730 | 88.25 | 734 |
| *P. sanctifelicis* | Gap | 1 | 1080 | 18 | 98.33 | 33 | 96.94 | 51 |
| | | 2 | 1080 | 22 | 97.96 | 64 | 94.07 | 86 |
| | | 3 | 1080 | 28 | 97.4 | 38 | 96.5 | 66 |

| | | | | | | | | |
|---|---|---|---|---|---|---|---|---|
| | | 4 | 359 | 2 | 99.44 | 2 | 99.44 | 4 |
| | | 5 | 1222 | 47 | 96.15 | 262 | 78.56 | 309 |
| | | 7 | 3508 | 72 | 97.95 | 92 | 97.38 | 164 |
| | | 8 | 1649 | 66 | 95.99 | 44 | 97.33 | 110 |
| | | 11 | 4676 | 902 | 80.71 | 737 | 84.24 | 1639 |
| *P.spD* | Gap | 1 | 6711 | 614 | 90.85 | 89 | 85.5 | 703 |
| | | 2 | 4320 | 95 | 97.8 | 80 | 98.15 | 175 |
| | | 3 | 360 | 38 | 89.44 | 126 | 65 | 164 |
| *P. sublinateum* | Forest | 1 | 360 | 0 | 97.22 | 10 | 97.22 | 360 |
| *P. umbricola* | Gap | 1 | 6256 | 123 | 98.03 | 87 | 98.61 | 210 |
| | | 2 | 2880 | 58 | 97.99 | 25 | 99.13 | 83 |
| Average | | | 4564.634146 | 217.4146341 | 95.1195122 | 406.725 | 87.5525 | 622.7560976 |
| Standard deviation | | | 3047.372522 | 342.4272898 | 5.37403524 | 450.8881257 | 15.2676035 | 710.3914193 |

**Table 4.** Number of each call type analyzed from acoustic data across 41 plants and 12 *Piper* species (see Methods).

| **Species** | **Habitat** | **Site** | **N files** | **Ins (FM-QCF)** | **Ins (FM-CF)** | **Feeding buzz** | **FM with harmonics** | **FM with harmonics (not Carollia)** | **FM without harmonics** | **Carollia calls** |
|---|---|---|---|---|---|---|---|---|---|---|
| *P. auritum* | Gap | 1 | 466 | 358 | 100 | 8 | | 2 | | |
| *P. colonense* | Forest | 1 | 400 | 1 | 4 | | | | | |
| | | 2 | 768 | 407 | 11 | 15 | | | 6 | |
| | | 3 | 1124 | 138 | 98 | 8 | | | 2 | |
| *P. cyanophyllum* | Forest | 1 | 210 | 54 | 19 | 5 | | 3 | | |
| *P. generalense (trigonum)* | Forest | 1 | 295 | 136 | 6 | 11 | | 2 | 3 | 2 |
| | | 2 | 296 | 104 | | 5 | | 3 | 9 | |
| | | 3 | 297 | 129 | 6 | 8 | | 3 | 4 | |
| | | 4 | 229 | 54 | | | 1 | | 1 | |
| | | 5 | 744 | 214 | 2 | 7 | | 1 | 1 | |
| | | 6 | 356 | | | | | | | |
| | | 7 | 701 | 333 | 21 | 14 | | 2 | | |

| | | | | | | | | | | |
|---|---|---|---|---|---|---|---|---|---|---|
| | | 8 | 210 | 51 | 24 | 3 | | | 4 | |
| *P. multiplinarium* | Gap | 1 | 1274 | 102 | 15 | 5 | | | | |
| *P. nudifolium* | Forest | 1 | 167 | 4 | 4 | | | 2 | | 1 |
| *P. pauliifolium (old: schiedeanum)* | Forest | 1 | 2237 | 307 | 49 | 10 | | 31 | 2 | |
| | | 2 | 3754 | 2240 | 116 | 396 | 34 | 17 | 298 | |
| | | 3 | 1100 | 311 | 48 | | | 4 | 3 | 2 |
| *P. reticulatum* | Forest | 1 | 239 | 28 | 26 | | | 8 | 1 | |
| | | 2 | 77 | 135 | | 6 | | | | |
| | | 3 | 976 | 176 | 2 | 3 | | 3 | | 2 |
| | | 4 | 856 | 134 | 42 | 8 | | 6 | 1 | |
| | | 6 | 744 | 83 | 38 | | | 2 | | |
| | | 7 | 1269 | 266 | 11 | 29 | | 15 | 13 | 3 |
| | | 8 | 789 | | | | | | | |
| | | 9 | 1097 | 314 | 16 | 24 | | 7 | 2 | |
| | | 10 | 734 | 198 | 42 | 10 | | | 427 | |

| | | | | | | | | | |
|---|---|---|---|---|---|---|---|---|---|
| *P. sanctifelicis* | Gap | 1 | 51 | 8 | | | | | |
| | | 2 | 86 | | 7 | | | | |
| | | 3 | 66 | 1 | 4 | 1 | | | |
| | | 4 | 4 | 1 | | | | | |
| | | 5 | 309 | 2 | 8 | | | | |
| | | 7 | 164 | 21 | 7 | 3 | | | 5 |
| | | 8 | 110 | | | | | | |
| | | 11 | 1639 | 240 | 162 | 18 | | 61 | 10 |
| *P. species D* | Gap | 1 | 703 | 686 | 74 | 11 | | 2 | 2 |
| | | 2 | 175 | 7 | 68 | 1 | | | |
| | | 3 | 164 | 37 | 62 | | | | |
| *P. sublinateum (biolleyi)* | Forest | 1 | 360 | 10 | 2 | | | | |
| *P. umbricola* | Gap | 1 | 210 | 16 | 37 | | | 1 | |
| | | 2 | 83 | 10 | 27 | | | | |

**Table S5.** Diet data compiled from the literature (Santana et al., 2021; Lopez & Vaughan, 2007; Maynard et al., 2019), where hundreds of fecal samples were collected and the percent of each *Piper* species in the fecal samples of each *Carollia* species (*C. sowelli, C. castanea, C. perspicillata*) were determined. We included the classification and habitat type for each *Piper* species.

| **Species** | **Habitat** | **Classification** | ***C. sowelli*** | ***C. castanea*** | ***C. perspicillata*** | **Max. %** | **Average %** |
|---|---|---|---|---|---|---|---|
| *P. aduncum* | open | mid-suc | 1.51898734 | 1.10132159 | 0.86580087 | 1.518987342 | 1.162036598 |
| *P. aequale (cabagranum)* | closed | late-suc | 0 | 0 | 0 | 0 | 0 |
| *P. prismaticum (augustum)* | closed | late-suc | 1.26582278 | 0.22026432 | 0.43290043 | 1.265822785 | 0.639662512 |
| *P. auritifolium* | closed | late-suc | 0.25316456 | 0 | 0 | 0.253164557 | 0.084388186 |
| *P. auritum* | open | early-suc | 15.1898734 | 0.88105727 | 12.987013 | 15.18987342 | 9.685981224 |
| *P. colonense* | open | mid-suc | 6.83544304 | 3.96475771 | 6.06060606 | 6.835443038 | 5.620268936 |
| *P. concepcionis* | closed | late-suc | 0.75949367 | 1.10132159 | 0.86580087 | 1.101321586 | 0.908872041 |
| *P. cyanophyllum (phytolaccifolium)* | closed | mid-suc | 0 | 0.22026432 | 0 | 0.220264317 | 0.073421439 |

| | | | | | | | |
|---|---|---|---|---|---|---|---|
| *P. darianense* | closed | mid-suc | 0 | 0 | 0 | 0 | 0 |
| *P. decurrens* | closed | mid-suc | 0 | 0 | 0 | 0 | 0 |
| *P. dryadum* | closed | mid-suc | 0 | 0 | 0 | 0 | 0 |
| *P. garagaranum* | closed | late-suc | 0 | 0 | 0 | 0 | 0 |
| *P. generalense* | closed | mid-suc | 0.25316456 | 1.54185022 | 3.8961039 | 3.896103896 | 1.897039558 |
| *P. holdridgeanum* | closed | late-suc | 0.25316456 | 0 | 0 | 0.253164557 | 0.084388186 |
| *P. evasum (imperiale)* | closed | late-suc | 0 | 0 | 0.43290043 | 0.432900433 | 0.144300144 |
| *P. melanocladum* | closed | late-suc | 0 | 0 | 0 | 0 | 0 |
| *P. multiplinervium* | open | early-suc | 7.08860759 | 18.2819383 | 18.6147186 | 18.61471861 | 14.66175485 |
| *P. nudifolium* | closed | mid-suc | 0 | 0.22026432 | 0 | 0.220264317 | 0.073421439 |
| *P. paulowniifolium* | closed | mid-suc | 0.50632911 | 2.42290749 | 0.43290043 | 2.422907489 | 1.120712345 |

| | | | | | | | |
|---|---|---|---|---|---|---|---|
| *P. peltatum* | open | early-suc | 0.50632911 | 0 | 0.86580087 | 0.865800866 | 0.45737666 |
| *P. pentagonum (cenocladum)* | closed | late-suc | 0 | 0 | 0 | 0 | 0 |
| *P. peracuminatum* | closed | mid-suc | 4.30379747 | 4.62555066 | 4.76190476 | 4.761904762 | 4.563750964 |
| *P. reticulatum* | open | mid-suc | 5.3164557 | 6.16740088 | 3.8961039 | 6.167400881 | 5.126653491 |
| *P. sanctifelicis* | open | early-suc | 35.443038 | 28.6343612 | 23.3766234 | 35.44303797 | 29.15134086 |
| *P. silvivagum* | closed | early-suc | 0.25316456 | 0.88105727 | 0.43290043 | 0.881057269 | 0.522374086 |
| *P. sp. D (hispidum)* | open | mid-suc | 2.53164557 | 3.08370044 | 5.19480519 | 5.194805195 | 3.603383735 |
| *P. sublineatum* | closed | mid-suc | 0.50632911 | 0 | 0 | 0.506329114 | 0.168776371 |
| *P. terrabanum* | closed | mid-suc | 0.25316456 | 0.44052863 | 0 | 0.440528634 | 0.231231064 |
| *P. tonduzii* | closed | late-suc | 0 | 0 | 0 | 0 | 0 |
| *P. umbricola* | open | early-suc | 7.5949367 | 11.4537445 | 6.49350649 | 11.453744 | 8.514062565 |

| | | | | | | | |
|---|---|---|---|---|---|---|---|
| | | | 1 | | | 49 | |
| *P. urophyllum* | closed | mid-suc | 0 | 0.66079295 | 0 | 0.660792952 | 0.220264317 |
| *P. urostachyum* | closed | late-suc | 0.75949367 | 0.44052863 | 0.43290043 | 0.759493671 | 0.544307579 |
| *P. xanthostachium* | closed | mid-suc | 0 | 0 | 0 | 0 | 0 |

**Tables 6a and 6b:** *Piper* species from the chemical dataset published in Santana et al. 2021, sorted by the concentration per species for each of the 15 most abundant chemicals among all *Piper* plants in the study. We included the habitat type (succession and forest/gap) as well.

| **Species** | **Succession** | **Habitat** | **N** | **Number of VOCs** | **Total emissions** | **Alpha caryophyllene** | **Germacrene D** | **Humulene** | **Beta pinene** | **Beta ocimene** | **Ethyl ester benzoic acid** |
|---|---|---|---|---|---|---|---|---|---|---|---|
| *P. aduncum* | mid | gap | 4 | 45 | 2149038.83 | 50771.50163 | 0 | 29129.33542 | 36700.01579 | 88865.06945 | 0 |
| *P. auritifolium* | late | forest | 3 | 30 | 54551.64943 | 6723.026706 | 188.5173518 | 10891.90165 | 0 | 0 | 0 |

| | | | | | | | | | | | |
|---|---|---|---|---|---|---|---|---|---|---|---|
| *P. colonense* | mid | gap | 7 | 53 | 336086.1168 | 13638.42508 | 88.56743684 | 14489.94391 | 15016.16142 | 0 | 0 |
| *P. concepcionis* | late | forest | 1 | 15 | 1012491.054 | 6691.716734 | 122027.89 | 0 | 0 | 67192.25845 | 24685.24902 |
| *P. darienense* | mid | forest | 2 | 16 | 91849.46052 | 1464.617769 | 0 | 0 | 0 | 0 | 1196.2236 |
| *P. evasum* | late | forest | 1 | 4 | 26157.16817 | 0 | 19647.07178 | 0 | 0 | 0 | 0 |
| *P. garagaranum* | late | forest | 1 | 6 | 358300.8222 | 70349.76018 | 0 | 64729.4162 | 0 | 167698.8488 | 0 |
| *P. generalense* | mid | forest | 5 | 30 | 260330.5138 | 16212.93424 | 139114.6478 | 11815.69602 | 0 | 2645.968872 | 1815.83632 |
| *P. multiplinervium* | early | gap | 4 | 24 | 278168.2472 | 89853.81223 | 37333.26804 | 1291.501196 | 21862.51607 | 0 | 13432.77695 |
| *P. nudifolium* | mid | forest | 2 | 24 | 103636.3404 | 56986.58672 | 0 | 5570.727955 | 0 | 664.440077 | 0 |
| *P. paulowniifolium* | mid | forest | 3 | 34 | 2040145.505 | 331420.8032 | 160401.2777 | 27031.66331 | 252901.8315 | 0 | 65289.07102 |
| *P. peltatum* | early | gap | 7 | 30 | 94475.3231 | 6743.054912 | 0 | 306.8400953 | 15529.16501 | 0 | 3304.495478 |
| *P. peracuminatum* | mid | forest | 6 | 54 | 77919.4 | 4753.208 | 9563.818 | 2518.3793 | 24410.341 | 7672.921 | 0 |

| | | | | | | | | | | | |
|---|---|---|---|---|---|---|---|---|---|---|---|
| | | | | | 4644 | 91 | 932 | 48 | 75 | 977 | |
| *P. prismaticum* | late | forest | 1 | 7 | 12864.52322 | 2560.994097 | 4213.65844 | 0 | 0 | 2061.807895 | 0 |
| *P. reticulatum* | mid | gap | 7 | 38 | 388150.5009 | 145054.2784 | 153706.9423 | 14124.20613 | 6289.491923 | 11345.24776 | 0 |
| *P. sanctifelicis* | early | gap | 34 | 104 | 332076.5465 | 13490.87042 | 2353.958133 | 1197.345968 | 4414.295557 | 3486.584744 | 1505.971868 |
| *P. silvivagum* | early | forest | 4 | 38 | 901877.6477 | 79779.8211 | 256428.7105 | 21961.05334 | 3498.384067 | 3076.920563 | 8479.863283 |
| *P. spD* | mid | gap | 1 | 14 | 186638.9391 | 54078.50359 | 44288.09822 | 4529.483438 | 27512.77739 | 1985.11003 | 2460.495715 |
| *P. sublineatum* | mid | forest | 2 | 19 | 121389.4916 | 2443.396942 | 8659.433355 | 0 | 4196.946122 | 0 | 0 |
| *P. umbricola* | early | gap | 2 | 23 | 579672.3874 | 90529.27032 | 249183.0545 | 12642.58031 | 34944.97138 | 18569.17689 | 35065.72281 |
| *P. urostachyum* | late | forest | 5 | 37 | 817541.9676 | 17663.47586 | 90049.41457 | 1920.305235 | 4895.160774 | 0 | 5522.610468 |

| **Species** | **Succession** | **Habitat** | **N** | **Alpha cubebene** | **Alpha phellandrene** | **p_Cymene** | **1-Dodecene** | **Beta elemene** | **3-methyl-2-** | **3-methyl-3-** | **Decanal** | **Beta myrcene** |
|---|---|---|---|---|---|---|---|---|---|---|---|---|

| | | | | | | | | | **undecene** | **undecene** | | |
|---|---|---|---|---|---|---|---|---|---|---|---|---|
| *P. aduncum* | mid | gap | 4 | 0 | 90186.91751 | 131884.5159 | 19022.09294 | 19495.3828 | 7933.616843 | 2992.273903 | 6066.075699 | 4101.439014 |
| *P. auritifolium* | late | forest | 3 | 534.818025 | 0 | 105.4621119 | 645.6321207 | 83.8550197 | 332.6397858 | 117.1614952 | 157.4943561 | 0 |
| *P. colonense* | mid | gap | 7 | 194.1385186 | 139964.3781 | 17509.65492 | 3787.228227 | 29271.40237 | 390.8158711 | 1517.285658 | 3262.210322 | 8446.167502 |
| *P. concepcionis* | late | forest | 1 | 0 | 7345.349486 | 6562.017794 | 0 | 0 | 0 | 0 | 0 | 11083.6206 |
| *P. darienense* | mid | forest | 2 | 0 | 0 | 0 | 0 | 0 | 0 | 0 | 0 | 0 |
| *P. evasum* | late | forest | 1 | 0 | 0 | 0 | 0 | 3070.261858 | 0 | 0 | 0 | 0 |
| *P. garagaranum* | late | forest | 1 | 22187.50171 | 0 | 0 | 0 | 0 | 0 | 0 | 0 | 0 |
| *P. generalense* | mid | forest | 5 | 1608.184152 | 0 | 0 | 0 | 2890.7712 | 0 | 0 | 0 | 0 |

| | | | | | | | | | | | | |
|---|---|---|---|---|---|---|---|---|---|---|---|---|
| *P. multiplinervium* | early | gap | 4 | 0 | 14631.04981 | 0 | 4892.057418 | 3358.014085 | 1855.746611 | 1852.004585 | 1829.406898 | 0 |
| *P. nudifolium* | mid | forest | 2 | 0 | 0 | 5068.430088 | 5963.091972 | 0 | 2282.931776 | 2253.292914 | 598.25717 | 0 |
| *P. paulowniifolium* | mid | forest | 3 | 490904.4425 | 18932.37911 | 69458.10093 | 0 | 0 | 0 | 0 | 0 | 10780.71302 |
| *P. peltatum* | early | gap | 7 | 1716.441984 | 302.1865024 | 8947.197177 | 0 | 0 | 0 | 0 | 0 | 112.2381208 |
| *P. peracuminatum* | mid | forest | 6 | 351.7438842 | 5114.927628 | 1271.053802 | 2528.011716 | 429.0736275 | 1569.781817 | 1024.090812 | 654.0145603 | 731.6862734 |
| *P. prismaticum* | late | forest | 1 | 0 | 0 | 0 | 0 | 0 | 0 | 0 | 0 | 0 |
| *P. reticulatum* | mid | gap | 7 | 4942.99677 | 1950.649869 | 0 | 4725.668849 | 0 | 2353.194329 | 1641.332771 | 1464.819504 | 0 |
| *P. sanctifelicis* | early | gap | 34 | 29447.09596 | 50316.11874 | 13395.30237 | 11352.69233 | 1732.807599 | 15062.26296 | 10314.81347 | 1600.923212 | 20.1365922 |
| *P. silvivagum* | early | forest | 4 | 20310.02841 | 907.4610614 | 0 | 0 | 0 | 0 | 0 | 0 | 2283.895428 |

| | | | | | | | | | | | | |
|---|---|---|---|---|---|---|---|---|---|---|---|---|
| *P. spD* | mid | gap | 1 | 4152.429928 | 0 | 11195.8953 | 0 | 18763.88233 | 0 | 0 | 0 | 1089.261061 |
| *P. sublineatum* | mid | forest | 2 | 0 | 0 | 0 | 33461.7148 | 0 | 11374.12393 | 7864.433078 | 2411.604225 | 0 |
| *P. umbricola* | early | gap | 2 | 7012.66521 | 0 | 0 | 19288.38708 | 3150.009926 | 7218.63605 | 4301.37228 | 13287.45834 | 6759.252714 |
| *P. urostachyum* | late | forest | 5 | 6799.869255 | 409428.2386 | 79098.06875 | 0 | 0 | 0 | 0 | 0 | 11518.93246 |
